# Direct access to millions of mutations by Whole Genome Sequencing of an oilseed rape mutant population

**DOI:** 10.1101/2022.08.17.504237

**Authors:** Srijan Jhingan, Avneesh Kumar, Hans-Joachim Harloff, Felix Dreyer, Amine Abbadi, Katrin Beckmann, Christian Obermeier, Christian Jung

**Affiliations:** Plant Breeding Institute, Christian-Albrechts-Kiel University, Olshausenstr. 40, 24098 Kiel, Germany; Department of Bioagricultural Sciences and Pest Management, Colorado State University, Fort Collins, Colorado, USA; NPZ Innovation GmbH, Hohenlieth-Hof, 24363 Holtsee, Germany; Department of Plant Breeding, Justus Liebig University Giessen, Heinrich-Buff-Ring 26-32, 35392 Giessen, Germany

**Keywords:** *Brassica napus*, ethyl methanesulfonate (EMS), EMS mutagenesis, TILLING, rapeseed

## Abstract

Induced mutations are an essential source of genetic variation in plant breeding. EMS mutagenesis has been frequently applied, and mutants have been detected by phenotypic or genotypic screening of large populations. In this study, a rapeseed M_2_ population was derived from M_1_ parent cultivar “Express” treated with EMS. Whole genomes were sequenced from fourfold (4x) pools of 1,988 M_2_ plants representing 497 M_2_ families. Detected mutations were not evenly distributed and displayed distinct patterns across the 19 chromosomes with lower mutation rates towards the ends. Mutation frequencies ranged from 32/Mb to 48/Mb. On average, 284,442 single nucleotide polymorphisms per M_2_ DNA pool were found resulting from EMS mutagenesis. 55% were C→T and G→A transitions, characteristic for EMS induced (‘canonical’) mutations, whereas the remaining SNPs were ‘non-canonical’ transitions (15%) or transversions (30%). Additionally, we detected 88,725 high confidence insertions and deletions (InDels) per pool. On average, each M_2_ plant carried 39,120 canonical mutations, corresponding to a frequency of one mutation per 23.6 kb. Roughly 82% of such mutations were located either 5 kb upstream or downstream (~56%) of gene coding regions or within intergenic regions (26%). The remaining 18% were located within regions coding for genes. All mutations detected by whole-genome sequencing could be verified by comparison with known mutations. Furthermore, all sequences are accessible via the online tool “EMS Brassica” (http://www.emsbrassica.plantbreeding.uni-kiel.de/), which enables direct identification of mutations in any target sequence. The sequence resource described here will further add value for functional gene studies in rapeseed breeding.

## Introduction

Oilseed rape (*Brassica napus* L.) is the primary oil crop in the world’s temperate regions and the third-largest seed oil and second-largest protein meal source globally (Wang et al., 2018). In 2019, ~34 million hectares yielded 70 million tons of oilseed rape globally (http://www.fao.org/faostat/). While the seeds are a major source of edible oil (45-50%), byproducts after oil extraction are conventionally utilized as animal feed and as a substrate for biodiesel production in Europe. Increasing and stabilizing the yield potential in combination with improved seed quality (e.g., reduced glucosinolate content) are the primary aims of rapeseed breeding.

Oilseed rape belongs to the family of crucifers (Brassicaceae). It is an allotetraploid (AACC, *2n* = 38) resulting from a spontaneous interspecific hybridization between the diploid AA (2n = 20) and CC (2n = 18) genomes of turnip rape (*Brassica rapa* L., syn. *campestris*) and cabbage (*Brassica oleracea* L.), respectively. Its origin was traced back to the Mediterranean region ~7,500 years ago and the total genome size is estimated to be ~1.13 Gigabases (Gb) (Chalhoub et al., 2014). Today, several high-quality rapeseed reference genomes are available. The “Darmor-bzh” was the first reference genome assembled using a European winter-type oilseed rape. The genome encompassed 314.2 Mb of the A sub-genome and 525.8 Mb of the C sub-genome, predicted with 101,040 gene models (Chalhoub et al., 2014). More recently, an improved long-read assembly of the “Darmor-*bzh*” reference genome (Rousseau-Gueutin et al., 2020) was published. Moreover, high-quality whole-genome assemblies of eight oilseed accessions across three ecotypes (Song et al., 2020), a winter-type (Lee et al., 2020) and a semi-winter (Chen et al., 2021) oilseed rape are now publically available.

Due to its short history of evolution and domestication, the genetic diversity within *B. napus* is low (Rahman, 2013). Ethyl methanesulfonate (EMS) induced mutagenesis has been used to create new allelic variation (Wang et al., 2008; Harloff et al., 2012; Gilchrist et al., 2013; Wells et al., 2014; Lee et al., 2018; Tang et al., 2020). In the past years, CRISPR-Cas technology to create targeted mutations has been successfully applied and numerous mutants have been published (Braatz et al., 2017; Karunarathna et al., 2020; Sashidhar et al., 2020; Zheng et al., 2020). Although targeted mutagenesis offers several advantages, random mutagenesis still has its importance in rapeseed breeding mainly because CRISPR-Cas mutants are legally classified as genetically modified organisms in the European Union and therefore their usage in practical breeding is limited (Jung & Till, 2021).

EMS mutant discovery in oilseed rape has conventionally relied on the activity of DNA mismatch specific endonucleases and polyacrylamide gel-based detection assays classically termed TILLING (Targeting Induced Local Lesions in Genomes) (Emrani et al., 2015; Braatz et al., 2018; Shah et al., 2018; Karunarathna et al., 2020) using pooled genomic DNA from M_2_ individuals of large mutant populations. Because this procedure is time-consuming and laborious, amplicon sequencing-based detection methods (Gilchrist et al., 2013; Wells et al., 2014; Sashidhar et al., 2019) gained increasing popularity owing to their efficiency and sensitivity. However, like conventional TILLING, detection of EMS mutations via amplicon sequencing approaches is restricted to single gene families with high sequence conservation.

Sequencing whole mutant populations, termed TILLING by sequencing (TbySeq), is the gold standard of mutant detection (Jung and Till, 2021). In a pioneering study, Krasileva et al. (2017) demonstrated a TILLING by exome sequencing approach to detect EMS-induced mutations in tetraploid and hexaploid wheat. The TbySeq approach has also been reported from several crops like rice (Abe et al., 2012), maize (Nie et al., 2021), tomato (Garcia et al., 2016), soybean (Lakhssassi et al., 2021), sunflower (Fanelli et al., 2021) and cotton (Fang et al., 2020). In a first TILLING by whole genome sequencing (TbyWGS) approach for oilseed rape, a limited number of EMS mutants were whole-genome sequenced to detect EMS-induced mutations (Tang et al., 2020).

This study aimed to develop a bioinformatic resource for detecting EMS-induced mutations on a genome-wide scale (Harloff et al., 2012). We sequenced the whole genomes of 1,988 M_2_ plants from an EMS mutagenized winter oilseed rape population. A TILLING by whole-genome sequencing (TbyWGS) pipeline was established which allows the identification of mutations within any genomic region of interest. The sequences can be screened via the online resource “EMS Brassica” (http://www.emsbrassica.plantbreeding.uni-kiel.de/). Thus, our TbyWGS platform constitutes a long-lasting sequence repository of mutants.

## Results

### Whole-genome sequencing reveals high mutation density

In order to gather first results on sequencing quality and mutant detection we first performed pilot experiments with fifty 4x pools sequenced at 10x coverage by mapping the raw reads to the Express617 v1 reference genome. The effective genome size of this reference assembly was 925,095,059 bp (Lee et al., 2020). After SNP detection and filtration, many SNPs were filtered into the “high confidence” category (data not shown) but later could not be validated via Sanger sequencing. Therefore, we decided to double the intended coverage to 20x for sequencing the 4x pools and sequenced pooled DNA from 1,988 M_2_ plants on the Illumina NovaSeq 6000 platform (Figure 1). The raw dataset encompassing a total of 497 4x pools represented an average of 35.8 gigabases (Gb) per pool. On average, ~116 million raw reads per pool were generated. In terms of read quality, Phred scores for all paired-end reads varied well above the optimum scores (Supplementary Figure 1). Mapping raw reads to the Express617 reference genome resulted in an average coverage depth of 34.2x.

**Figure 1.**
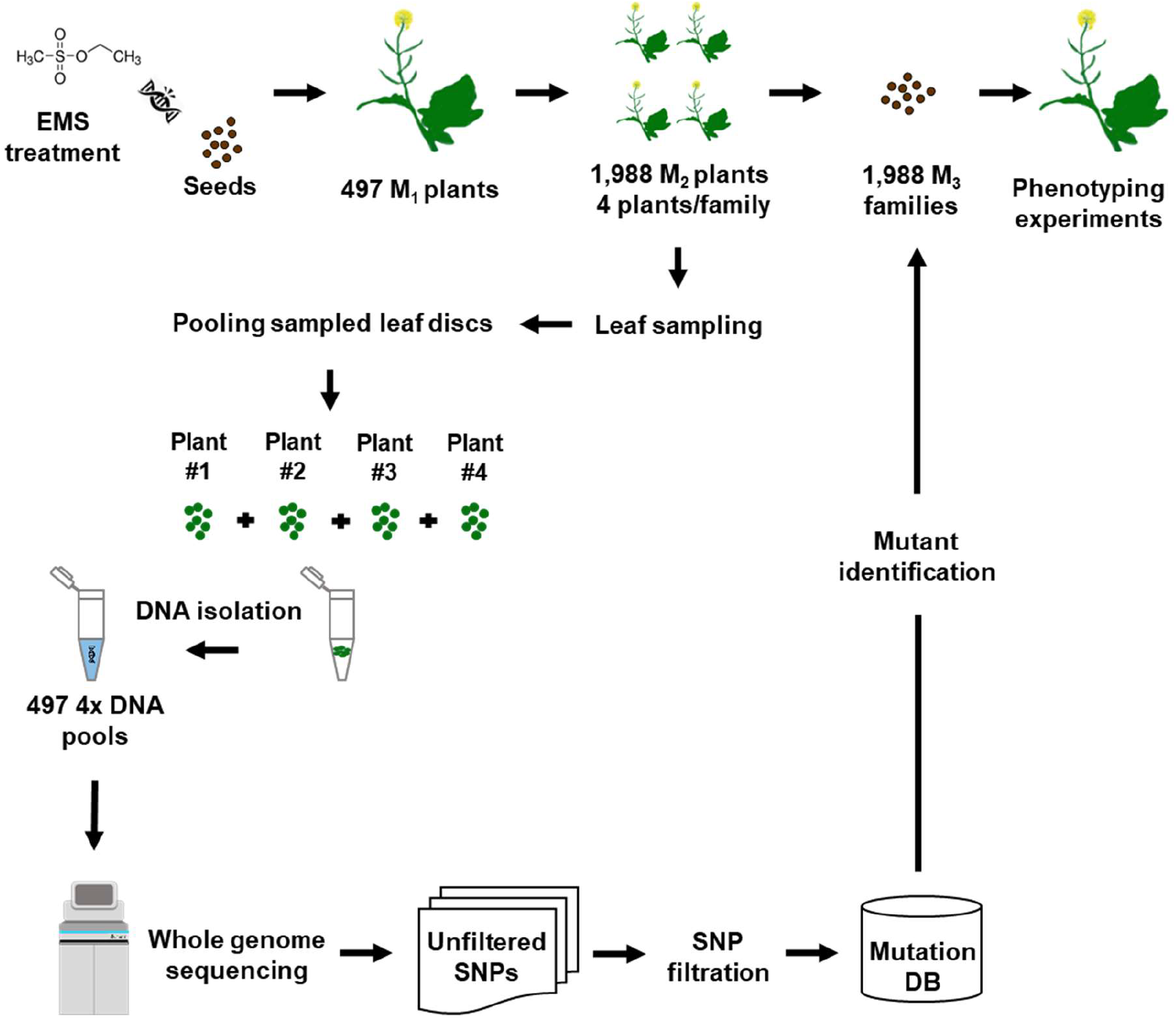
Workflow of the mutant detection pipeline. M_2_ seeds were harvested from 497 M_1_ plants and four plants/M_2_ family were grown. Leaf material was sampled from 1,988 M_2_ plants as leaf discs. Leaves from each family were pooled and DNA was isolated and sequenced on the Illumina NovaSeq 6000 platform. Raw reads were mapped to the Express617 reference genome. SNPs were called and filtered using GATK 4.1.4.1 and custom UNIX scripts. Raw SNPs were filtered based on canonical EMS mutations (C→T and G→A transitions) possessing read depth (DP) ≥10 allele depth (AD) =12.5%-60% and quality controlled for mapping quality (MQ) ≥30. Mutation effects were predicted using Ensembl Variant Effect Predictor (VEP) tool. High confidence mutations from each pool were merged into a single accessible database. After identification of desirable M_2_ mutants, corresponding M_3_ plants can be analyzed to verify for phenotypic effects.

We then calculated the number and frequency of SNPs. Variant calling revealed 843,000 unfiltered SNPs per pool on average. Using our SNP filtering criteria (read depth ≥10, allele depth =12.5%-60% and mapping quality ≥30), on average, 156,480 SNPs per pool were C→T and G→A transitions, characteristic for EMS induced mutations (Table 1). This corresponds to an average of 39,120 EMS type mutations per single M_2_ plant. Following the terminology used by Fanelli et al. (2021), we termed the EMS type C→T and G→A mutations as “canonical” transitions and all others as “non-canonical” transitions or transversions. We termed them collectively as single nucleotide variants (SNVs). We detected an average of one C→T and G→A mutation per 23.6 kb of the genomic sequence of the Express617 reference genome (925 Mb). Noteworthy, the 45% share of non-canonical transitions (15%) or transversions (30%) was almost equal to that of the canonical C→T and G→A mutations among the high confidence SNPs (Figure 2). Additionally, on average, we detected 88,725 high confidence insertions and deletions (InDels) per M_2_ DNA pool. However, we restricted subsequent sequence analyses to SNVs only.

**Figure 2.**
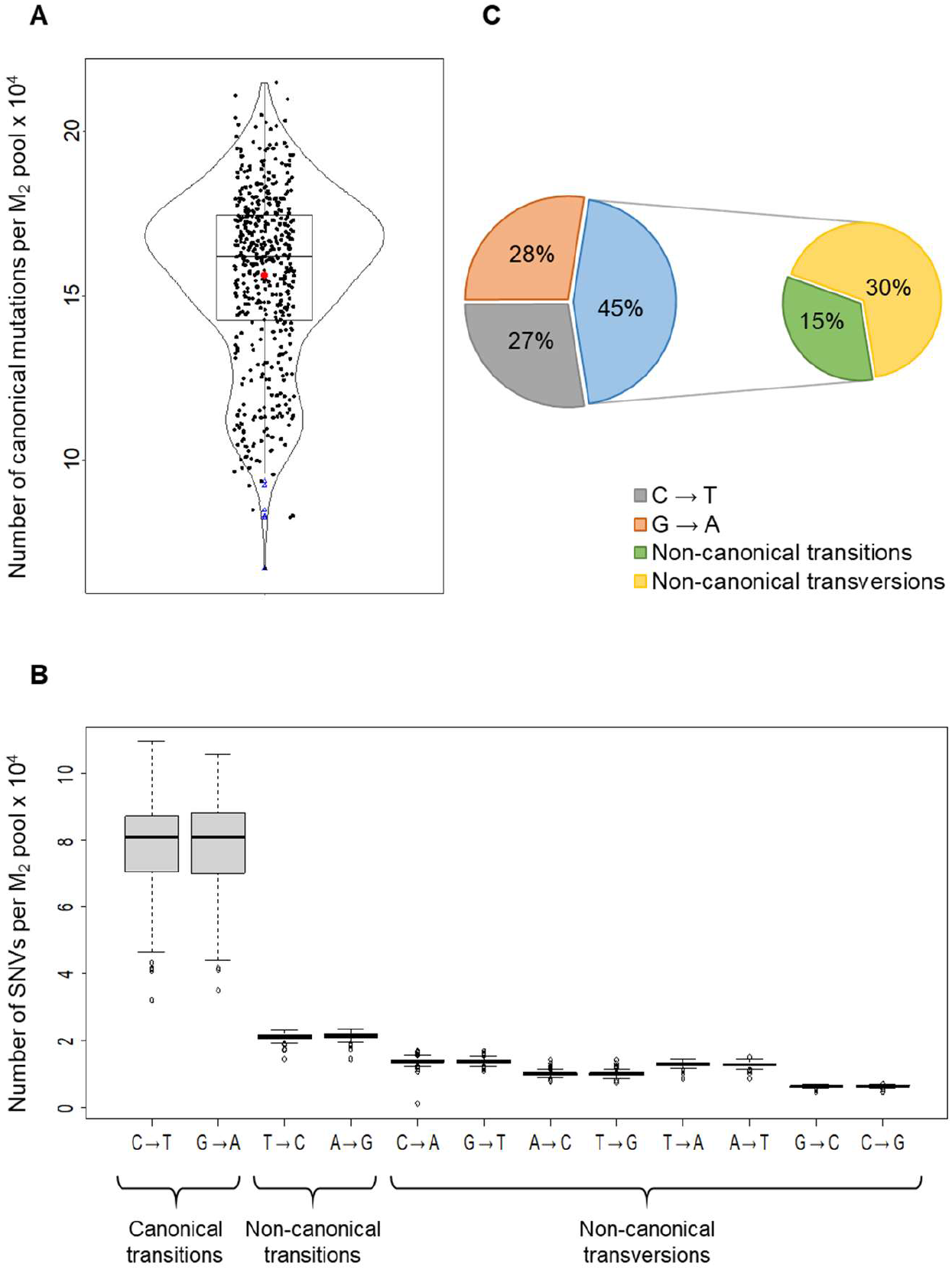
Analysis of SNP counts from 1,988 sequenced M_2_ plants constituting 497 4x pools. (A) Violin plot showing the number of canonical C→T and G →A transitions. Black dots and blue triangles depict the number of filtered SNPs from each of the sequenced pools and outliers (pools with a very low number of called SNPs), respectively. The red dot represents the mean of SNP counts from all pools. (B) Boxplots showing the number of individual SNP types observed and (C) Percentage share of single nucleotide variants (SNVs) by type. C→T and G→A transitions accounting for 55% of total filtered SNVs were named as “canonical” mutations. All other nucleotide substitutions (45%) have been named as “non-canonical” transitions or transversions. All SNVs were filtered based on SNP type, minimum read depth (DP) ≥10, allele depth (AD) =12.5%-60% and mapping quality (MQ) ≥30 parameters. All boxplots show the upper and lower quartiles separated by the median (horizontal line). Whiskers represent maximum and minimum values.

**Table 1.** Summary statistics of SNPs detected after whole genome sequencing (including regions without chromosomal annotations) of 1,988 M_2_ rapeseed plants assembled in 497 4x DNA pools. SNPs were filtered based on DP≥10, AD=12.5%-60% and MQ≥30. Characteristic C→T and G→A transitions were named as “canonical EMS-type” transitions. All other substitutions were named as “non-canonical” transitions or transversions.

|  | Canonical transitions |  | Non-canonical transitions |  | Transversions |  |  |  |  |  |  |  |
| --- | --- | --- | --- | --- | --- | --- | --- | --- | --- | --- | --- | --- |
| SNP | C → T | G → A | T → C | A → G | C → A | G → T | A → C | T → G | T → A | A → T | G → C | C → G |
| Number of SNPs <sup>[a]</sup> | 78,170 | 78,310 | 21,073 | 21,174 | 13,770 | 13,776 | 9,942 | 9,930 | 12,839 | 12,754 | 6,290 | 6,414 |
| Sum | 156,480 |  | 42,247 |  | 85,715 |  |  |  |  |  |  |  |
| % | 27.0 | 28.0 | 7.0 | 7.0 | 5.0 | 5.0 | 3.0 | 3.0 | 5.0 | 4.0 | 2.0 | 2.0 |

[a] Average of 497 4x pools analyzed DP: Read depth, AD: Allelic depth, MQ: Mapping quality

### Detecting functional mutations

We questioned how many high confidence EMS mutations could have a putative effect on gene function. First, we analyzed mutations within predicted gene models from the Express617 reference. We reasoned that an SNV might alter the function of the encoded protein if it 1) introduces a premature stop codon, 2) results in a splice variant, 3) confers a non-synonymous amino acid substitution, or 4) causes a change in the translation start site. On average, 12,129 mutations per M_2_ DNA pool fulfilled these criteria (Table 2). 75% of all characterized mutations were located within annotated regions of the Express617 reference genome (chromosomes A01-A10 and C01-C09), whereas 25% were located on non-annotated scaffolds (occupying ~17% of the reference genome) that could not be confidently anchored to any chromosome (Supplementary Table 1). Out of all high confidence canonical mutations located within annotated chromosomes, 0.4%, 7.0%, 4.1%, and 4.9% were predicted as nonsense, missense, synonymous and intronic, respectively (Table 2). Out of all canonical mutations located within coding regions, 2.3% were nonsense mutations, 40.4% were missense mutations, 24.1% were synonymous mutations and 28.7% were located within introns (Supplementary Figure 2). Start site loss and splice site variants were observed as the rarest type of mutations contributing <1% of the total EMS-type mutations (Figure 3).

**Figure 3.**
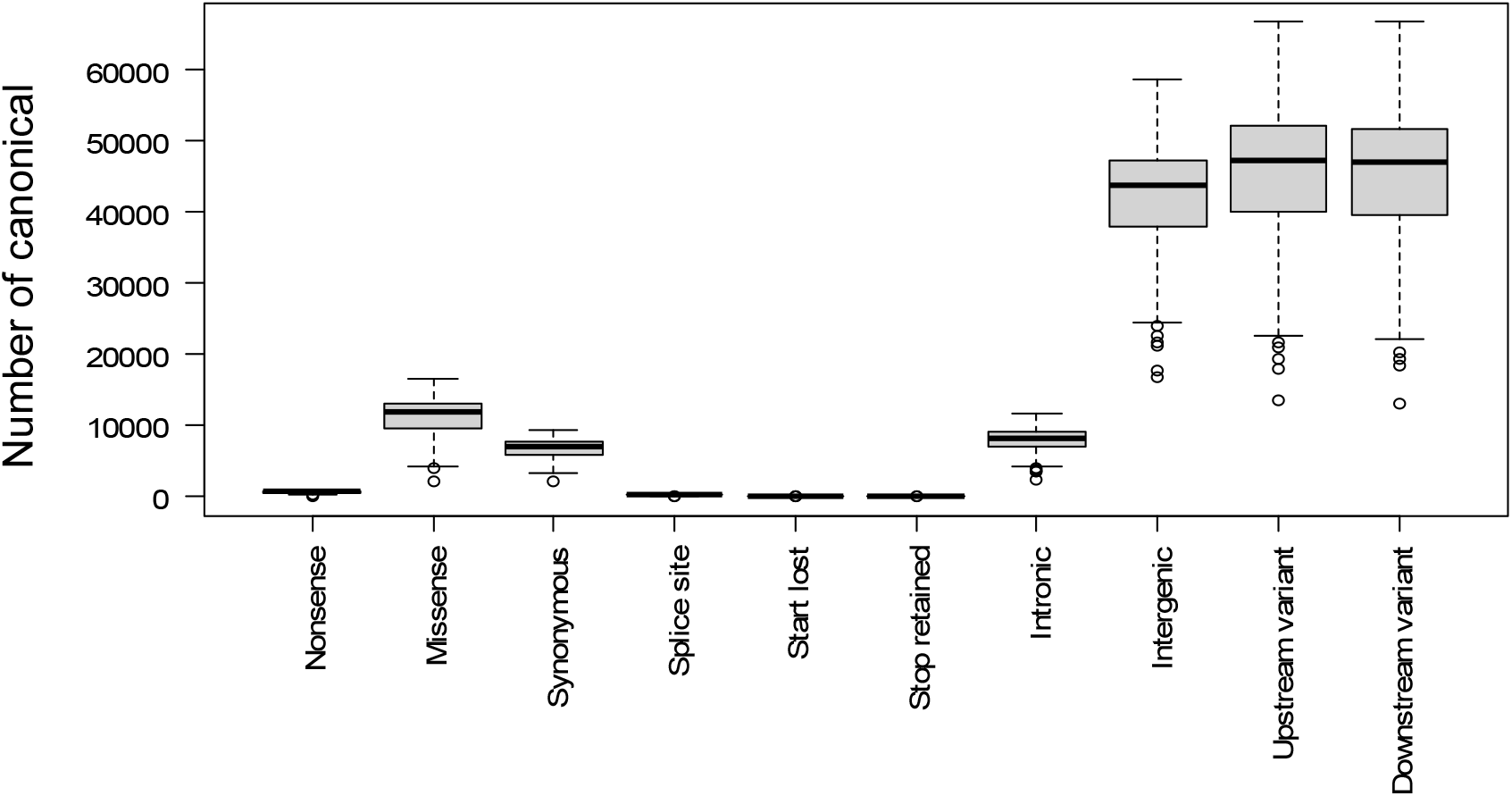
Characterization of C→T and G →A mutations with predicted effects on a genome-wide scale. Boxplots show distribution of SNP effects from the sequenced 4x pools. The Upper and lower quartiles are separated by the median (horizontal line). Whiskers represent maximum and minimum values. Circles depict outliers. Mutation effects are predicted using SNPs filtered with DP>10, AD=12.5%-60% and MQ≥30 parameters. The Ensembl Variant Effect Predictor release 99 was used in offline mode. Mutation effects were characterized within all predicted gene models using the general feature format (GFF) file of the Express617 reference genome. Splice site variants include acceptor and donor site mutations. Stop retained mutations refer to unchanged STOP codons in spite of induced mutation(s). Upstream and downstream variants are located within a distance of 5 kb from the transcription START and STOP sites, respectively. DP: Read depth, AD: Allele depth and MQ: Mapping quality.

**Table 2.**
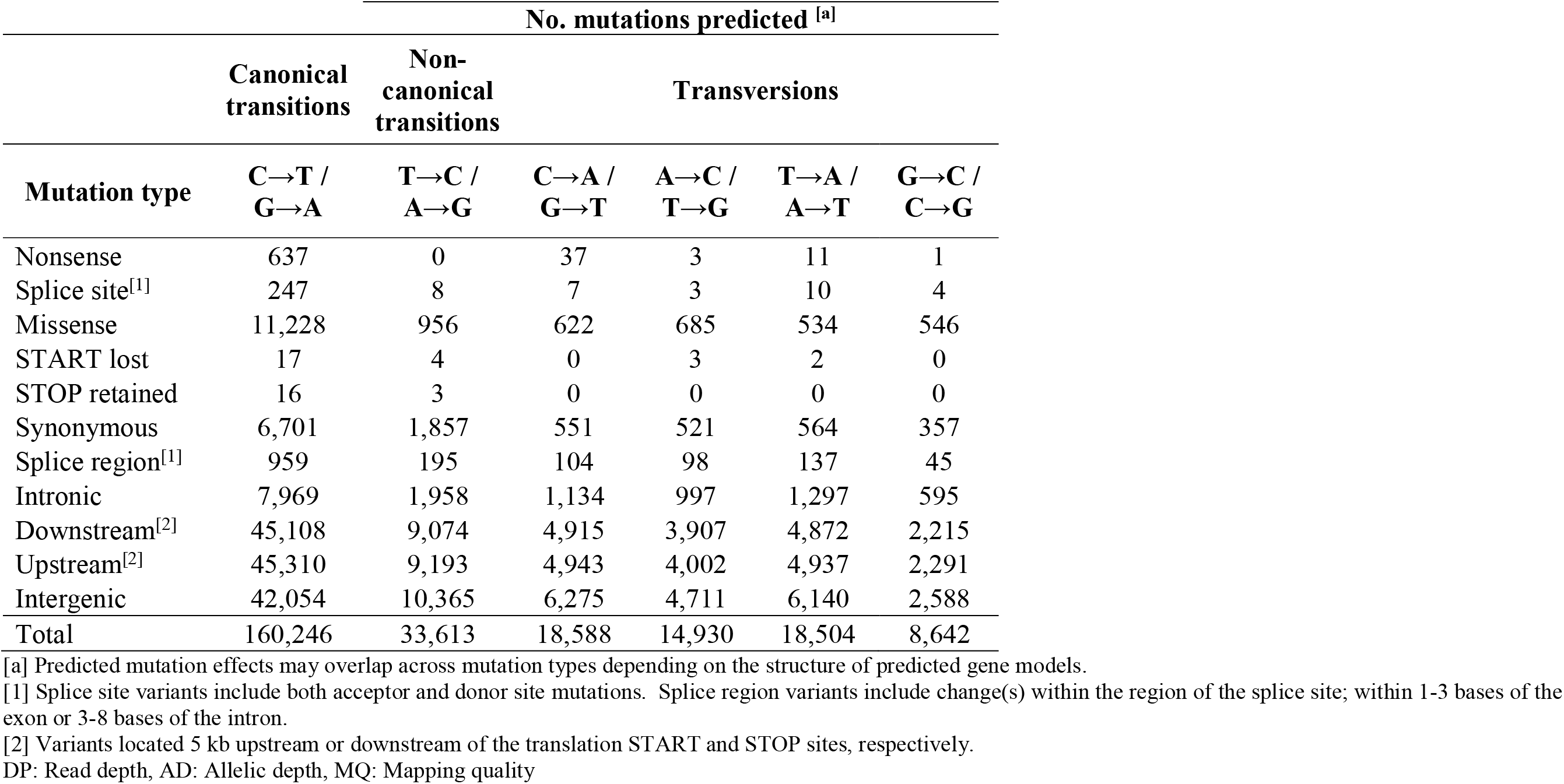
Summary statistics of predicted mutation effects within the predicted gene models of the Express617 genome. Only “EMS-type” C→T and G→A transitions from 1,988 sequenced M_2_ plants were used. Mutation effects are predicted using filtered SNPs with DP≥10, AD=12.5%-60% and MQ≥30 from chromosomally annotated regions (chrA01-A10 and chrC01-C09). The Ensembl Variant Effect Predictor release 99 was used in offline mode. All gene IDs were extracted from the general feature format (GFF) of the Express617 reference genome (Lee et al., 2020).

| Mutation type | No. mutations predicted <sup>[a]</sup> |  |  |  |  |  |
| --- | --- | --- | --- | --- | --- | --- |
|  | Canonical transitions | Non-canonical transitions | Transversions |  |  |  |
|  | C→T /<br>G→A | T→C /<br>A→G | C→A /<br>G→T | A→C /<br>T→G | T→A /<br>A→T | G→C /<br>C→G |
| Nonsense | 637 | 0 | 37 | 3 | 11 | 1 |
| Splice site<sup>[1]</sup> | 247 | 8 | 7 | 3 | 10 | 4 |
| Missense | 11,228 | 956 | 622 | 685 | 534 | 546 |
| START lost | 17 | 4 | 0 | 3 | 2 | 0 |
| STOP retained | 16 | 3 | 0 | 0 | 0 | 0 |
| Synonymous | 6,701 | 1,857 | 551 | 521 | 564 | 357 |
| Splice region<sup>[1]</sup> | 959 | 195 | 104 | 98 | 137 | 45 |
| Intronic | 7,969 | 1,958 | 1,134 | 997 | 1,297 | 595 |
| Downstream<sup>[2]</sup> | 45,108 | 9,074 | 4,915 | 3,907 | 4,872 | 2,215 |
| Upstream<sup>[2]</sup> | 45,310 | 9,193 | 4,943 | 4,002 | 4,937 | 2,291 |
| Intergenic | 42,054 | 10,365 | 6,275 | 4,711 | 6,140 | 2,588 |
| Total | 160,246 | 33,613 | 18,588 | 14,930 | 18,504 | 8,642 |

[a] Predicted mutation effects may overlap across mutation types depending on the structure of predicted gene models. [1] Splice site variants include both acceptor and donor site mutations. Splice region variants include change(s) within the region of the splice site; within 1-3 bases of the exon or 3-8 bases of the intron. [2] Variants located 5 kb upstream or downstream of the translation START and STOP sites, respectively. DP: Read depth, AD: Allelic depth, MQ: Mapping quality

We then analyzed the functional effects of mutations other than C→T and G→A transitions (Supplementary Figure 3). Roughly 85% of such mutations were located either 5 kb upstream or downstream (~27% each) of gene coding regions or within intergenic regions (31%). The remaining 15% were located within regions coding for genes (Supplementary Figure 4). As predicted, 3.6% of these mutations could have drastic effects in the form of nonsense (0.05%), missense (3.5%), and splice site (0.03%), or even as START site (0.01%) mutations. On the contrary, 10.4% do not confer functional effects since they were characterized as silent mutations like synonymous (4%) or intronic (6.3%) and rare mutations where the stop codon is retained (0.003%) (Table 2).

### Sequence analyses reveal patterns of mutation frequency and distribution

Our mutation detection approach operates on the whole-genome scale. Therefore, we investigated the distribution of EMS mutations across all chromosomes, including intergenic regions and 5 kb upstream and downstream regions. To check for a possible bias in mutation frequencies for the A and C sub-genomes of oilseed rape, we calculated the number and density of high confidence canonical mutations across all annotated chromosomes (Table 3). As expected, the number of C sub-genome mutations (76,818) was significantly higher than A sub-genome mutations (41,412) because in the assembled genome the C genome exceeds the A genome by 173.25 Mb (Figure 4). Surprisingly, mutation frequencies on average were higher for the C sub-genome (41 mutations/Mb) than the A sub-genome (35 mutations/Mb).

**Figure 4.**
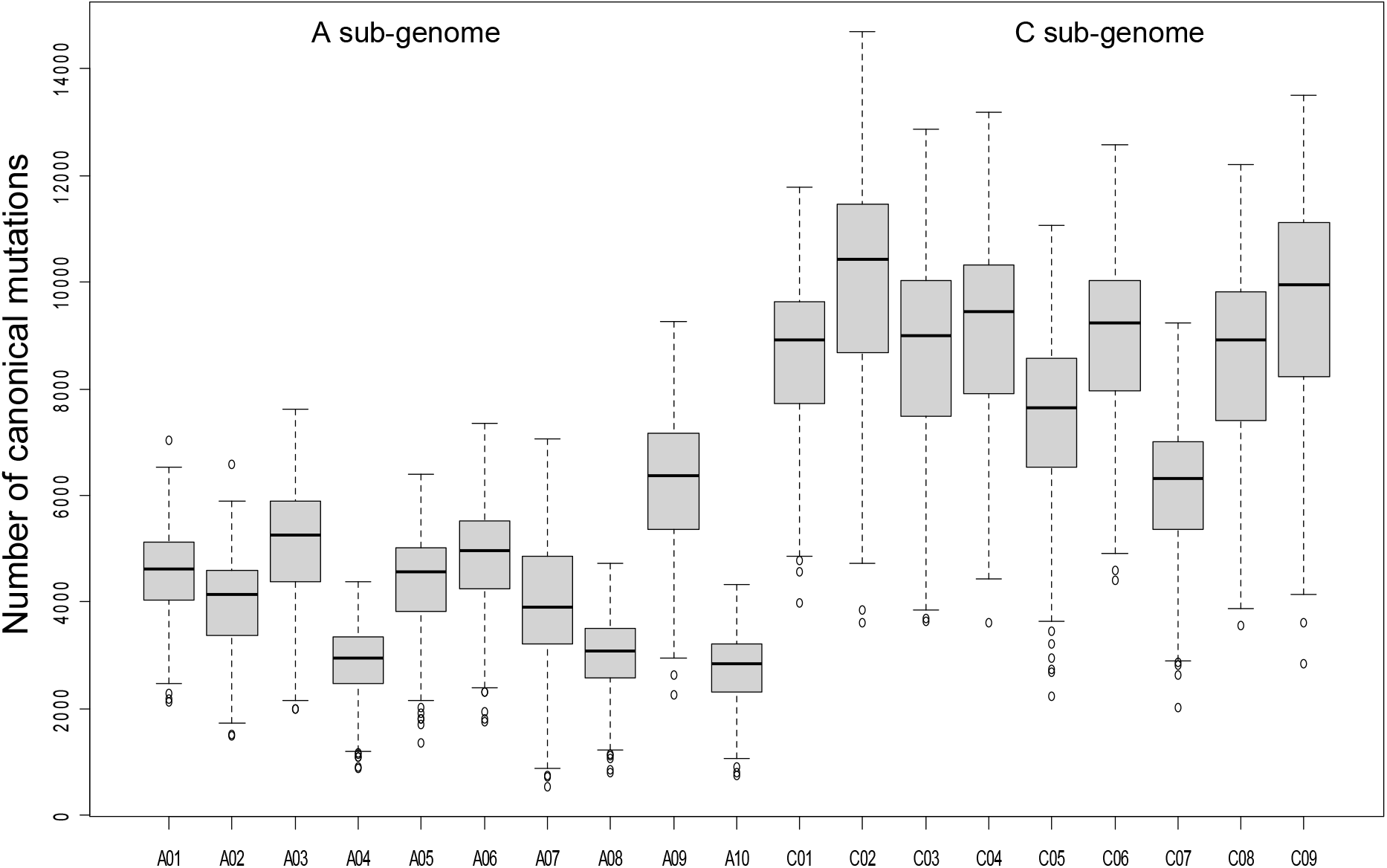
Number of EMS-induced transitions across the A- and C sub-genomes of 497 sequenced M_2_ pools. Boxplots represent the average number of SNPs originating from the regions of the Express617 genome annotated with chromosomes (chrA01 - A10 and chrC01 - C09). Black circles represent individual data points for SNP counts per sequenced 4x pool. The Upper and lower quartiles are separated by the median (horizontal line). Whiskers represent maximum and minimum values. Circles depict outliers.

**Table 3.**
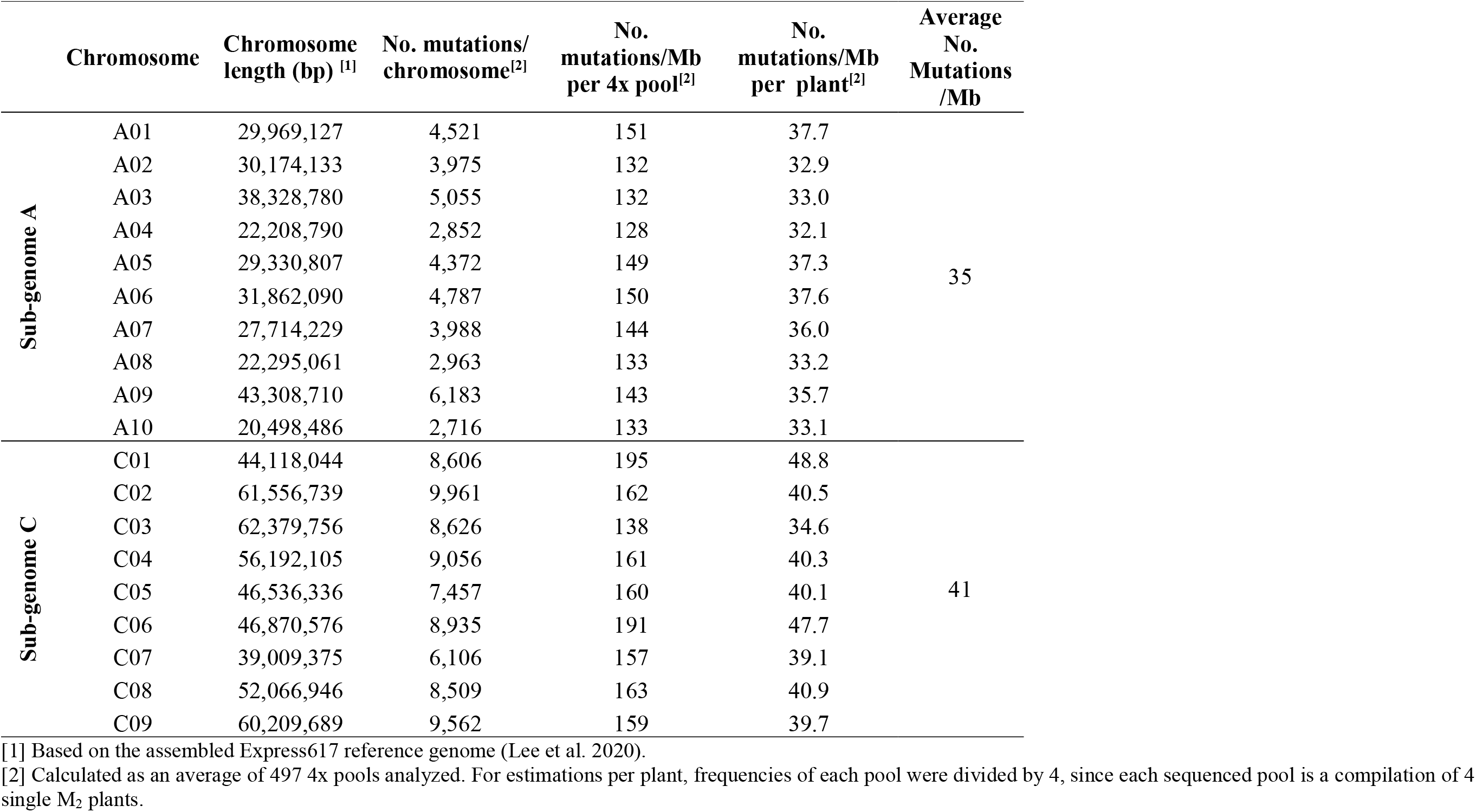
Summary statistics of EMS type transitions in the A and C sub-genomes of the Express617 mutant population. Mutation counts and frequencies were calculated for individual chromosomes of the two sub-genomes (chrA01-A10 and chrC01-C09) for all sequenced 4x pools.

|  | Chromosome | Chromosome length (bp) <sup>[1]</sup> | No. mutations/<br>chromosome<sup>[2]</sup> | No.<br>mutations/Mb<br>per 4x pool<sup>[2]</sup> | No.<br>mutations/Mb<br>per plant<sup>[2]</sup> | Average<br>No.<br>Mutations<br>/Mb |
| --- | --- | --- | --- | --- | --- | --- |
| Sub-genome A | A01 | 29,969,127 | 4,521 | 151 | 37.7 | 35 |
|  | A02 | 30,174,133 | 3,975 | 132 | 32.9 |  |
|  | A03 | 38,328,780 | 5,055 | 132 | 33.0 |  |
|  | A04 | 22,208,790 | 2,852 | 128 | 32.1 |  |
|  | A05 | 29,330,807 | 4,372 | 149 | 37.3 |  |
|  | A06 | 31,862,090 | 4,787 | 150 | 37.6 |  |
|  | A07 | 27,714,229 | 3,988 | 144 | 36.0 |  |
|  | A08 | 22,295,061 | 2,963 | 133 | 33.2 |  |
|  | A09 | 43,308,710 | 6,183 | 143 | 35.7 |  |
|  | A10 | 20,498,486 | 2,716 | 133 | 33.1 |  |
| Sub-genome C | C01 | 44,118,044 | 8,606 | 195 | 48.8 | 41 |
|  | C02 | 61,556,739 | 9,961 | 162 | 40.5 |  |
|  | C03 | 62,379,756 | 8,626 | 138 | 34.6 |  |
|  | C04 | 56,192,105 | 9,056 | 161 | 40.3 |  |
|  | C05 | 46,536,336 | 7,457 | 160 | 40.1 |  |
|  | C06 | 46,870,576 | 8,935 | 191 | 47.7 |  |
|  | C07 | 39,009,375 | 6,106 | 157 | 39.1 |  |
|  | C08 | 52,066,946 | 8,509 | 163 | 40.9 |  |
|  | C09 | 60,209,689 | 9,562 | 159 | 39.7 |  |

[1] Based on the assembled Express617 reference genome (Lee et al. 2020). [2] Calculated as an average of 497 4x pools analyzed. For estimations per plant, frequencies of each pool were divided by 4, since each sequenced pool is a compilation of 4 single M<sub>2</sub> plants.

In the next step, we searched both sub-genomes for mutation hotspots by visualizing EMS mutations in 1 Mb non-overlapping windows (Figure 5). While canonical mutations were evenly distributed across all chromosomes in general, we found regions with significantly increased or decreased mutation densities (Figure 5). As a general tendency, mutation frequencies were lower towards the end of chromosomes except for A06, C03 and C06, where mutation frequencies were higher in the telomeric regions.

**Figure 5.**
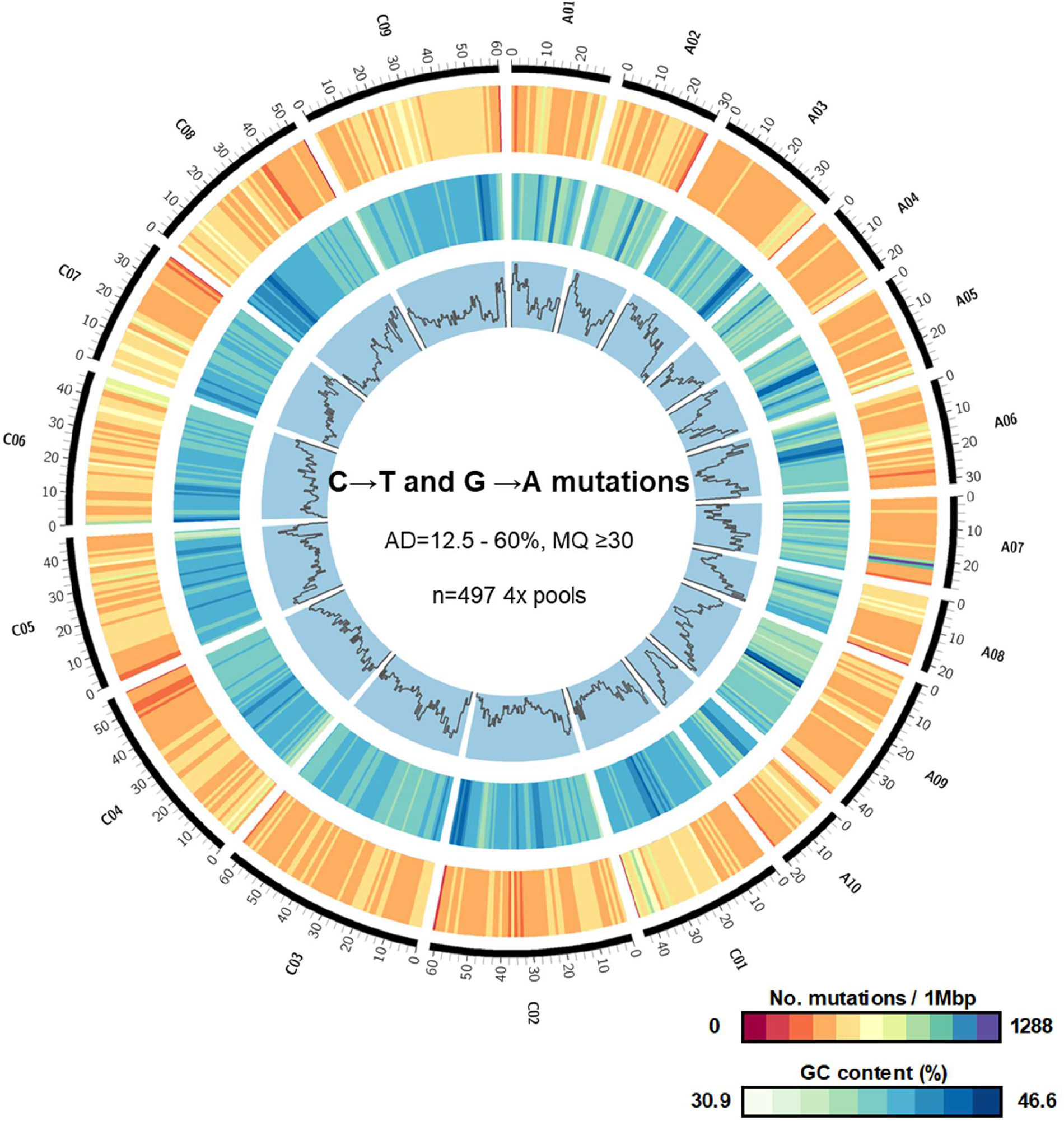
Visualization of C→T and G →A mutations at the whole genome-scale. The Circos plot shows four tracks (outside to inside) representing chromosomes A01-A10 and C01-C09 (black), number of mutations (dark red to violet), and GC content (%) per 1 Mb windows (white to dark blue) and regions coding for genes. Chromosomal lengths are in Megabases (Mb). Mutation densities were calculated per 1 Mb windows across all chromosomes from all sequenced 4x pools. Regions coding for genes were calculated as the number of base pairs per 1 Mb that were annotated within gene models. Mutation densities and GC content were plotted using “scale_log_base = 0.5” values. Non-overlapping windows were used for calculations.

Furthermore, we compared the distribution of canonical and non-canonical mutations for all chromosomes. For this, we used all C→T and G→A transitions and then all other mutation types that qualified as high confidence SNPs (read depth ≥10, allele depth =12.5%-60%, and mapping quality ≥30) as distinct sets. Interestingly, a similar distribution of high confidence canonical and non-canonical mutations per 1 Mb non-overlapping windows was observed across most chromosomes (Supplementary Figure 5). This suggests that the distribution of the non-canonical mutations is not random across chromosomes but follows a defined distribution pattern like that observed for the C→T and G→A transitions.

### Validation of functional mutations

The functional mutations detected by the TbyWGS approach were verified in two ways. We chose four gene families controlling agronomically important traits in oilseed rape. First, we verified mutations in the four gene families previously detected by a conventional gel-based TILLING of the Express617 TILLING population (Harloff et al., 2012; Emrani et al., 2015; Karunarathna et al., 2020). *SFAR* (*SEED FATTY ACID REDUCER*) genes encode GDSL lipases which control a wide range of primary and secondary functions in plants. Plants possessing EMS-induced functional mutations in *BnSFAR* genes had displayed an elevated seed oil content (Karunarathna et al., 2020). *REF1* and *SGT* genes encode for enzymes UDP-glucose:sinapic acid glucosyltransferase and sinapaldehyde dehydrogenase/coniferaldehyde dehydrogenase, respectively. Both genes play a crucial role in the biosynthesis of sinapine, an important anti-nutritive compound in oilseed rape. EMS-induced loss of function mutations had been detected in both genes. Double and triple mutants showed a significant reduction of the seed sinapine content (Harloff et al., 2012; Emrani et al., 2015). To verify the mutations by our TbyWGS approach, we identified 7 M_2_ families with the respective M_2_ mutants. We then screened whole genome sequences of the 4x pools harboring these M_2_ families (Supplementary Table 2). As a result, all seven mutations could be found, proving the reliability of our TbyWGS pipeline (Supplementary Figure 6).

As a next step, we screened the WGS dataset for new mutations. We chose five genes involved in the glucosinolate biosynthesis and transportation pathway. *MYB28* and *CYP79F1* genes encode for an R2-R3-MYB transcription factor (Gigolashvili et al., 2008) and a cytochrome P450 enzyme (Reintanz et al., 2001), respectively. Both genes are involved in the core structure formation of aliphatic glucosinolates in oilseed rape. The *GTR (GLUCOSINOLATE TRANSPORTER*) genes have been characterized by seed-specific glucosinolate transportation activity in *Brassica* species (Nour-Eldin et al., 2017). We detected 12 M_2_ DNA pools with nonsense mutations for all selected gene families (Supplementary Figure 7). Then, we isolated genomic DNA from each of the 48 M_2_ plants. DNA fragments were amplified by PCR using paralog-specific primers flanking the putative mutation sites (Supplementary Table 3). Sanger sequencing revealed a success rate of 100% because all PCR fragments contained the expected mutations (Supplementary Figure 8).

### A web-based interface for screening the mutant population

Our TbyWGS resource “EMSBrassica” provides access to 78,150,284 high confidence canonical EMS mutations. We developed a web-based resource enabling the user-friendly and convenient detection of high confidence C→T and G→A mutations with predicted effects for ease of screening. Access will be given via the institute’s web page. As a first step, the Express617 reference genome assembly and the supplemented GFF file assembled by Lee et al. (2020) must be downloaded from https://doi.org/10.5281/zenodo.3524259. By using BLAST queries, genomic regions of interest within this reference assembly concerning the chromosome and their exact locations will be identified. Then, the WGS database can be searched based on the “Chromosome”, “Start” and “End” information. By screening the database, mutations within genomic regions of interest can be identified from the respective 4x pools. The database “EMS Brassica” is accessible at http://www.emsbrassica.plantbreeding.uni-kiel.de/.

## Discussion

This study aimed to develop a database of oilseed rape mutant sequences for identifying putative functional mutations on a genome-wide scale. Based on the available Express617 mutant population, the reference genome Express617 (Lee et al., 2020) and whole genome sequencing of M_2_-Pools of mutants, we have established a TILLING by whole genome sequencing (TbyWGS) protocol to detect mutations in any sequence of the respective rapeseed mutant genome. M_2_ plants with the desired mutation can be directly identified to perform functional studies in the M_3_ offspring. This procedure avoids the laborious gel-based mutant screening and the long-term storage of high-quality M_2_ DNA needed for conventional TILLING.

In our TbyWGS approach, we used a 4x pooling strategy instead of the original 8x pooling previously used for gel-based screening of the Express617 mutant population (Harloff et al., 2012). We found that low sequence coverage results in the false identification of sequencing errors or artifacts as genuine mutations. Therefore, a 20x sequence coverage was chosen, which theoretically results in five raw reads from each M_2_ plant. The degree of heterozygosity in the Express617 inbred line (F11) used for EMS mutagenesis is expected to be below 0.0005%. Therefore, we expect that >99.9995 % of the SNPs detected within our M_2_ families are caused by EMS mutagenesis and not by residual heterozygosity. We adjusted the SNP filtering criteria to remove low-quality SNPs with poor mapping quality and sequencing errors. According to classical Mendelian genetics, the frequency of mutant alleles in a segregating M_2_ generation is expected to be 50% if we assume the M_1_ to be hemizygous for the mutation. However, this value might be lower due to a mosaic of mutated and non-mutated cells which is expected for almost all M_1_ plants. Also, poor fitness of mutant gametophytes and low vitality of homozygous M_2_ plants can account for decreased mutant frequencies in the M_2_ offspring. In addition, we can expect a statistical bias caused by the low number of M_2_ plants in each pool since only four M_2_ plants per pool were selected. In the worst case, only one plant out of four might carry a mutant allele. In another case, the mutation might be more frequent due to an increased representation of mutated plants carrying one or more mutant alleles within an M_2_ pool. Respecting the Mendelian segregation and biases due to mosaic M_1_ and fitness of M_2_ plants in all sequenced pools, it was reasonable in our screening to select a lower limit for the allelic depth of 12.5% to cover rare mutations in each pool. On the other hand, the upper limit from the expected Mendelian average of 50% to 60% was increased to correct for an overrepresentation of mutations in a given M_2_ pool. A recommended mapping quality (MQ) ≥30 was consistently ensured for all called variants for quality control. Moreover, we only used SNPs from loci that had a total coverage depth of at least ten mapped reads at that position. SNV calling was purposely based on these relaxed parameters to avoid false-negative classification of putative useful mutants in target genes based on too strict filtering parameters. This might also have resulted in the calling of a high percentage of non-canonical SNVs. However, validation by Sanger sequencing of known mutants revelead that the positive mutant calls were all correctly called and thus make this data set a useful resource for identification of genotypes with mutations in selected target genes.

Mutation frequencies can vary between different rapeseed EMS mutant populations. (Wang et al., 2008; Harloff et al., 2012; Gilchrist et al., 2013; Wells et al., 2014; Tang et al., 2020). This can be due to several factors like EMS concentration and permeability and the physiology and developmental stage of the treated tissue (Henry et al., 2014). Based on previous studies using conventional gel-based assays, the mutation frequency in our EMS population ranged between 1/12 kb to 1/72 kb (Harloff et al., 2012; Guo et al., 2014; Emrani et al., 2015; Braatz et al., 2018; Shah et al., 2018; Karunarathna et al., 2020). While using an amplicon sequencing approach, a mutation frequency of 1/27 kb was reported from the same TILLING platform by Sashidhar et al. (2019). These differences can be explained because mutation frequencies were estimated based on the length of the amplicons analyzed. Moreover, only protein-coding sequences were studied. In our TbyWGS approach, we estimated a mutation frequency of one high confidence EMS mutation per 23.6 kb of the Express617 reference genome, which is in line with previous estimations from the same EMS population. However, it is expected that mutation frequencies calculated exclusively for gene coding regions will vary when compared with the distribution of mutations for the entire genome. Therefore, a more suitable and direct comparison can be drawn from the mutation frequencies of 1/23.64 kb to 1/303.86 kb observed by Tang et al. (2020) from 20 whole genome sequenced single oilseed rape EMS mutants. Mutation frequencies in EMS resources from other crops like rice (Till et al., 2007), tomato (Garcia et al., 2016) (Wells et al., 2014) and wheat (Krasileva et al., 2017) were estimated to be 1/530 kb, 1/125 kb and as high as 30-40 per kb, respectively.

The non-random distribution of mutation across the whole genome is an important result of our study. For the first time, we report a lower density of EMS-induced mutations at the ends of the chromosomes (Figure 5). Using the representative group of 19 annotated chromosomes in the Express617 reference assembly (chromosomes A01-A10 and C01-C09), we calculated mutation frequencies from different regions of the genome (Table 3, Supplementary Figure 5). While the higher number of mutations in the C sub-genome is explained by its genome size, the striking difference in the mutation frequencies between the two sub-genomes deserves a further explanation. One reason could be different GC contents. In the assembled Express617 genome, the average GC contents per 1 Mb windows of the A and C sub-genomes are 35.2% and 36.2%, respectively. Although this difference is not high, we reason that a high GC content invariably affects mutation frequencies since mutations observed were primarily C→T and G→A transitions (55%). Therefore, regions with high GC content could be potential mutation hotspots. Furthermore, it can be speculated that both sub-genomes are not randomly targeted by EMS. At the time of EMS treatment, the vast majority of cells are in the interphase. It is known from studies with polyploid species e.g. wheat (Fussell & Moens, 1987; Martinez-Perez et al., 2001) that sub-genomes adopt the ‘Rabl configuration’ and are located separately in the interphase nucleus. This could have an impact on their accessibility to EMS resulting in varying mutation densities. Also, the frequency and distribution of repetitive elements alter the conformation of the chromatin fibre during the interphase. It is tempting to speculate that EMS targets DNA differentially in condensed and relaxed chromatin.

EMS is believed to confer mainly C→T and G→A transitions (Till et al., 2006). This is reflected in our TbyWGS resource since 55% of all high confidence SNPs were canonical mutations. Using *in-vitro* DNA ethylation experiments, Sega (1984) has demonstrated a range of possible modifications other than C→T and G→A transitions resulting from DNA depurination and depyrimidation. It was observed that even transversions could originate since a random nucleotide was inserted opposite apurinic or apyrimidic sites. Since we observed 45% non-canonical mutations in our TbyWGS approach, we questioned why such mutations were rarely observed in our previous studies with the Express617 mutant population. Traditionally, functional genomic studies utilizing EMS mutants have focused on identifying mutations within coding regions (Emrani et al., 2015; Braatz et al., 2018; Karunarathna et al., 2020). Moreover, these studies used conventional detection assays utilizing the CEL I endonuclease, an S1 single strand-specific nuclease that cleaves heteroduplexes of mutant and wild-type DNA. It has been demonstrated that although CEL I identifies all sequence variants, including InDels (Oleykowski et al., 1998), it has the highest preference for cleaving heteroduplexes arising from G→C, G→A, G→T and C→G mutations (Oleykowski et al., 1998; Yang et al., 2000; Triques et al., 2007). Using our TbyWGS approach, we observed that for high confidence SNPs located in gene coding regions, 76.3% were within this group of mutations highly preferred by CEL I. Out of this 66.7% were canonical mutations and only 9.6% were G→C, G→T or C→G mutations. We reason that such non-canonical mutations were rarely reported from previous conventional screenings mainly due to two reasons. Firstly, mutant detections were done mainly within gene coding regions. Secondly, most non-canonical transitions and transversions within gene coding regions represented the least preferred mismatch sites for CEL I cleavage (Oleykowski et al., 1998). Within gene coding regions, a low frequency of non-canonical mismatch heteroduplexes favored by CEL I is possibly a reason for low detection rates in the past.

We detected 45% non-canonical transitions and transversions. This is in line with a recent report where >50% non-canonical transitions, transversions and InDels were observed in three out of ten whole genome sequenced M_2_ EMS mutagenized rapeseed plants (Tang et al. (2020). Moreover, our observation is in line with several reports utilizing conventional and sequencing-based detection methods for EMS mutants from rice (Till et al., 2007), maize (Lu et al., 2018), barley (Caldwell et al., 2004), soybean (Lakhssassi et al., 2020) and sunflower (Fanelli et al., 2021). In these studies, the share of non-canonical transitions and transversions ranged from 12.3% to 31% (Krasileva et al., 2017).

Presently, our mutant database encompasses the canonical transitions. Adding a screening method for non-canonical transitions, transversions could be an interesting option and a possible expansion in the future, especially regarding the regulatory 5 kb up-and downstream regions of genes. Deletions and insertions are expected to be present in our M_2_ population due to EMS treatment, an effect that has been reported before in a study on DNA ethylation experiments (Sega, 1984). Moreover, via an exome sequencing approach Krasileva et al. (2017) have confirmed the presence of small InDels (<20 bp) in EMS mutagenized populations of tetraploid and hexaploid wheat. While small (<10 bp) InDels can be detected in our Illumina sequenced population, screening for large InDels requires long-range sequencing technology.

In conclusion, our web-accessible whole-genome sequencing mutant platform is an unprecedented resource that can serve as a strong foundation for molecular breeding and functional genomics in oilseed rape research. The process of mutant identification will now altogether bypass cumbersome detection methods in a time and cost-effective manner. Since the resource represents whole-genome sequencing data, it not only enables functional characterization of EMS mutations within genes but also offers a novel opportunity to analyze mutations within regulatory sequences. In this respect, investigation of mutations detected upstream and downstream of genes or even within intergenic regions is now a possibility and perhaps a compelling proposition for researchers and plant breeders alike. Although CRISPR-Cas mediated editing of multiple gene families has gained immense momentum in the past years, especially for the improvement of oilseed rape (Braatz et al., 2017; Karunarathna et al., 2020; Sashidhar et al., 2020; Zheng et al., 2020), legal restrictions within the European Union, have deterred application in current breeding programs. In this regard, EMS mutagenized functional mutants hold great promise for successful integration in plant breeding and crop improvement (Jung & Till, 2021).

## Summary

This study presents a novel TILLING by whole-genome sequencing (TbyWGS) platform to detect EMS-induced mutations on the whole-genome level from a winter rapeseed EMS mutant population. The resource contains 78,150,284 high confidence EMS-induced C→T and G→A mutations originating from whole-genome sequencing data from 1,988 M_2_ plants. On average, each plant possessed ~39,000 mutations with a frequency of 1/23.6 kb on the genome level. Roughly 82% of the mutations were located in 5 kb upstream or downstream (~28% each) of gene coding regions or within intergenic regions (26%). The remaining 18% were located within regions coding for genes. Of these, 0.4%, 7.0%, 4.1%, and 4.9% were predicted as nonsense, missense, synonymous and intronic mutations, respectively. The study also notes distinct distribution patterns of EMS induced mutations on a genome-wide scale. Our designed web-based resource incorporates whole-genome sequencing data enabling the user-friendly detection of EMS mutations in any genomic region of interest. This effectively bypasses the need for cumbersome gel-based mutation screening techniques for detection of EMS induced functional mutations in oilseed rape.

## Methods

### Plant material

The offspring of the winter oilseed rape inbred line Express617 (F11) had been treated with EMS and M_2_ and M_3_ generations had been produced (Harloff et al., 2012). 1,988 M_2_ plants (4 plants/family) representing 497 M_2_ families (4 plants/family) were grown under greenhouse conditions (22°C, 16 h light) at the NPZ Innovation GmbH. Plants were vernalized for 12 weeks at 4°C. All M_2_ plants were self-pollinated and M_3_ seeds were harvested.

### DNA isolation and whole-genome sequencing

Twenty 4 mm disks were taken from leaves of each of the M_2_ plants before vernalization. DNA was isolated from bulked leaf samples of each M_2_ family. The pooled leaf samples were lyophilized (ALPHA 1-4 LDplus, Martin Christ Gefriertrocknungsanlagen GmbH, Germany) for 96 h. Genomic DNA of 497 4x DNA pools was isolated using the DNeasy Plant Mini Kit following the manufacturer’s protocol (QIAGEN GmbH, Germany) and sequenced on an Illumina NovaSeq 6000 platform using 150bp paired ends reads and a 350bp insert DNA library with 20x depth of coverage (Novogene, Co., Ltd., China).

### Data processing and SNP detection

Raw data were obtained in the FASTQ format. The quality of raw reads was checked using the FastQC v0.11.5 (Andrews, 2010) and MultiQC (Ewels et al., 2016) programs. Based on the quality checks, samples representing low sequencing outputs or reads possessing poor Phred scores, deviating GC count per read, and a high number of ambiguously called bases were removed from the analysis. A long-read genome assembly of Express617 was used as the reference genome (Lee et al., 2020). Raw reads from all 4x pools were mapped to the Express617 oilseed rape reference, using BWA-MEM (Li, 2013) with default parameters for local alignment. SAMtools was used to convert, sort, and index the resulting files to binary alignment map files. The average depth of coverage per pool was measured using the SAMtools –depth tool.

Using Picard tools AddOrReplaceReadGroups and MarkDuplicates, mapped reads were pre-processed before variant calling. Variants were called using the GATK 4.1.4.1 HaplotyeCaller program (Poplin et al., 2018). The resulting Variant Call Format (vcf) files were filtered for single nucleotide polymorphisms characteristic to the canonical EMS type C→T and G→A transitions. SNPs were retained based on read depth (DP) ≥10, mapping quality (MQ) ≥30, and allele depth (AD) between 12.5% - 60% parameters using bcftools and custom UNIX scripts. Allele depth of mutations was calculated as the percentage share of reads possessing mutations from total reads (read depth, DP) mapped at that position. These SNPs are hereafter referred to as “high confidence” SNPs. The same criteria were used to create a subset of variants representing all other nucleotide substitutions referred to as “non-canonical” transitions and transversions. The Variant Effect Predictor (VEP) tool (McLaren et al., 2016) was used in the offline mode to characterize and predict SNP effects on the polypeptide level. The General Feature Format (GFF) file for the Express617 reference assembly (Lee et al., 2020) representing predicted protein models was used to characterize all SNP effects on a genome-wide scale.

### Estimation of mutation frequencies

To approximate the mutation frequency per kb (kilobases) of the reference genome, the number of high confidence SNPs along the chromosomal length for each of the annotated chromosomes (A01-A10 and C01-C09) and the non-annotated share of the reference genome was calculated. A genome-wide mutation density was calculated by dividing the number of mutations by the effective genome size (~925 Mb). Within 1 Mb (megabases) non-overlapping windows, the frequency of C→T and G→A transitions and all other transitions and transversions were then calculated from each of the 4x pools for all chromosomes. A moving average for the number of SNPs across all sequenced pools per 1 Mb non-overlapping windows was calculated for all chromosomes. The General Additive Model (GAM) in the ggplot2 package on R (Wickham, 2009) was used to smoothen the curves while plotting values. The number of putative functional mutations conferred by C→T and G→A transitions and all other mutations were then calculated.

### Developing the web-based mutant database

The web-accessible database EMSBrassica (http://emsbrassica.plantbreeding.uni-kiel.de/) was built using Django (https://www.djangoproject.com/), a Python-based free and open-source web framework. PostgreSQL (http://www.postgresql.org), an open-source object-relational database system, was used as the database engine. Webpages were built using HTML and Bootstrap (https://getbootstrap.com/).

## Declarations

### Availability of data and materials

The datasets supporting the conclusions of this article are available in the NCBI Sequence Read Archive - BioProject: PRJNA758762 (https://dataview.ncbi.nlm.nih.gov/object/PRJNA758762?reviewer=pmlp61jqoo028afrj45the0ull). All individual files representing SNP effects from whole genome sequenced pools are accessible at https://doi.org/10.5281/zenodo.6617711. All codes used can be made available from the authors upon request.

### Authors’ contributions

SJ, HJH, AA, FD, CO and CJ designed the research. AA, FD and KB generated the plant material and developed the fourfold pools. SJ conducted the experiments and analyzed the data. SJ and AK designed and developed the webpage. HJH and CJ supervised the research. SJ wrote the original draft. HJH, AA, CO and CJ reviewed and edited the manuscript. All authors participated in the discussion and revision of the manuscript. The authors read and approved the final manuscript.

## Acknowledgements

We thank the Institute of Clinical Molecular Biology in Kiel for providing Sanger sequencing as supported in part by the DFG Clusters of Excellence “Precision Medicine in Chronic Inflammation” and “ROOTS”.

## Supplementary data

### Supplementary Tables

**Supplementary Table 1.**
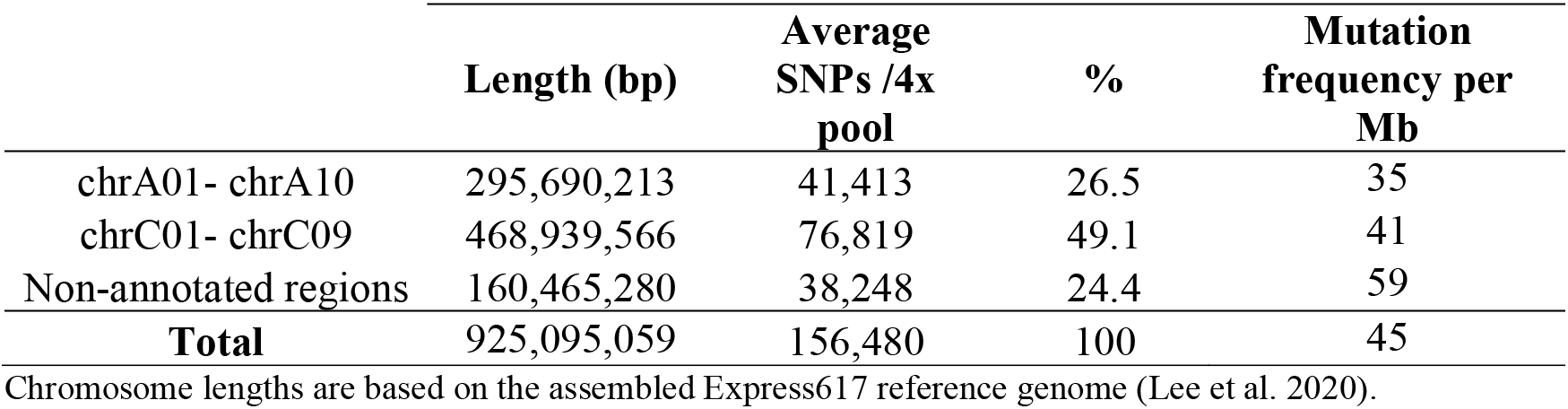
Summary of EMS-type transitions originating from the annotated and non-annotated regions of the Express617 genome. Regions of the Express617 reference assembly without chromosomal annotations for the A- or C sub-genomes are denoted as non-annotated regions. Share of C→T and G→A transitions located within annotated and non-annotated regions was calculated as an average from 497 sequenced 4x pools.

**Supplementary Table 2.**
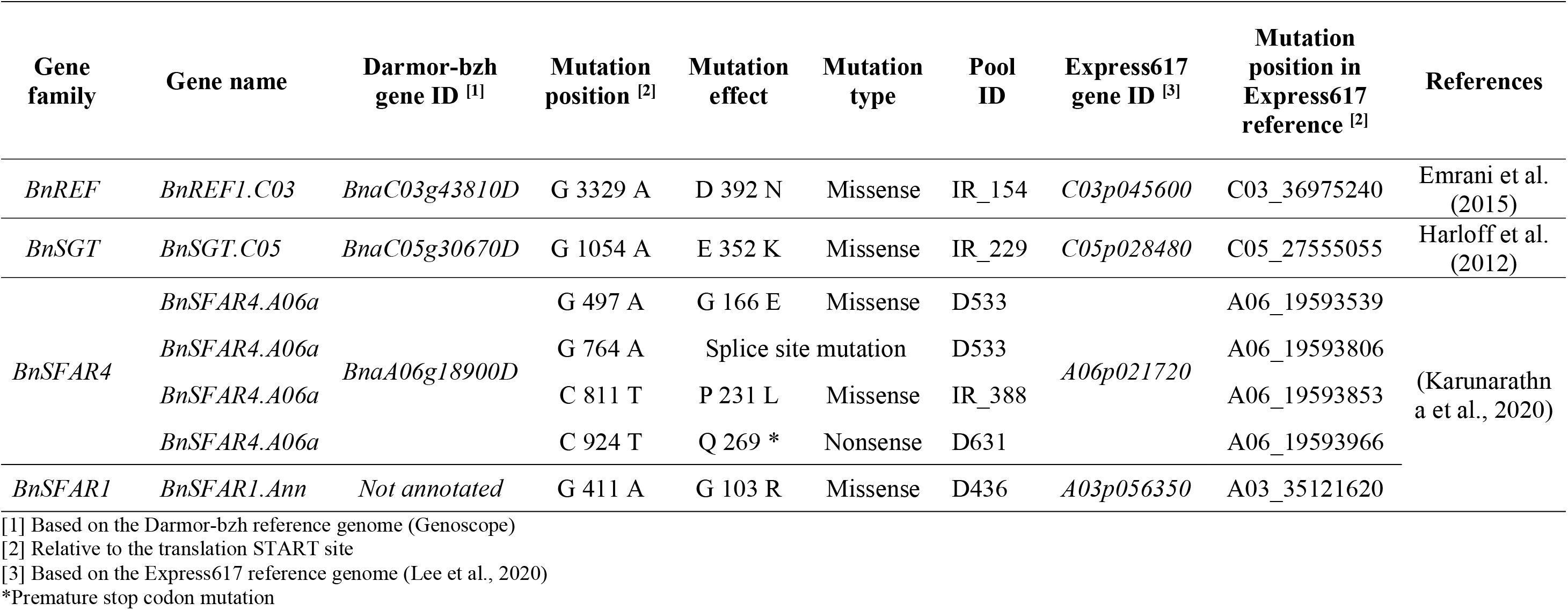
Summary of validation experiments to confirm mutations in 7 M_2_ DNA pools harboring previously detected and characterized EMS mutations within 4 candidate gene families, *BnREF, BnSGT, BnSFAR4* and *BnSFAR1*. For each of the selected mutants, corresponding M_2_ families were identified. Individual read alignments from each of the 4x pools (named with prefix “D” or “IR”) representing the selected mutant M_2_ families were visualized for regions harboring expected mutations.

**Supplementary Table 3.**
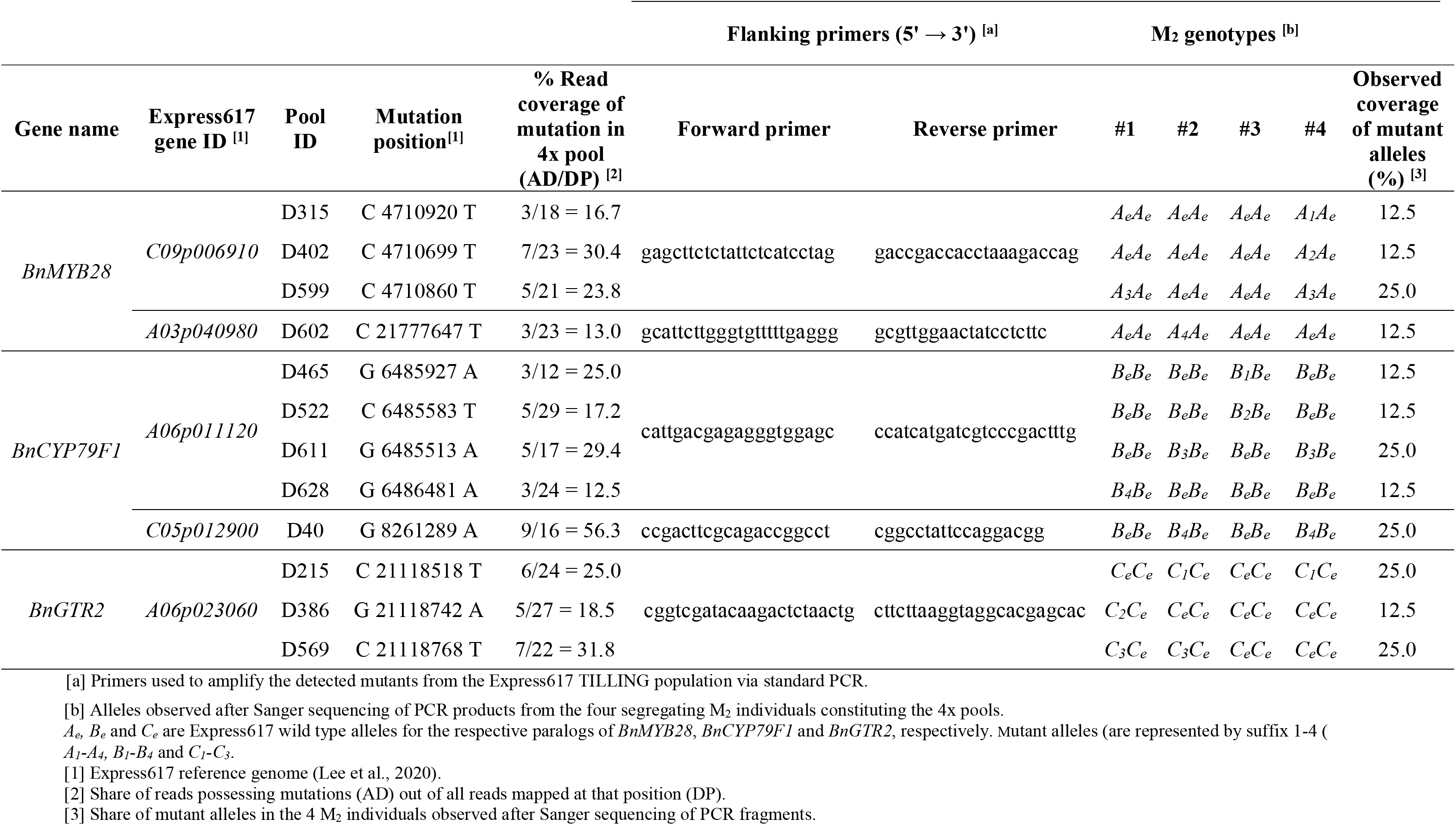
Summary of validation experiments to confirm the presence of mutations in 12 pools harboring EMS mutations within candidate gene families *BnMYB28, BnCYP79F1* and *BnGTR2*. 4x pools (named with prefix “D”) with nonsense mutations for the candidate genes were first identified. Genomic DNA was isolated separately from the leaf samples of the 4 individuals (#1-4) bulked in each pool. Standard PCR with locus-specific primers was used to amplify regions encompassing the detected mutations. PCR fragments were Sanger sequenced to validate mutation presence.

### Supplementary Figures

**Supplementary Figure 1** - Screenshot of the FastQC reports showing a graphical overview of quality checks for raw reads from all sequenced 4x pools. Mean quality scores represent Phred scores for individual bases called across the length 150 bp paired-end reads. Phred scores for all 4x pools are within the optimum levels of >30 (blocks marked in green). Raw reads with Phred score = 20-26 are tolerated (yellow zones), Phred score ≤20 (red zones) are rejected. Analysis was done via FastQC v0.11.5 and MultiQC.

**Supplementary Figure 2** - Share of EMS-induced mutations predicted to have putative functional effects (A) on a genome-wide scale and (B) specifically within coding regions. SNP effects were first predicted on a genome-wide scale for all predicted gene models on the Express617 reference genome. The share of mutation effects within coding regions was calculated using SNPs that were present within and including the translation START and STOP sites. Splice site variants include the sum of splice acceptor and donor site mutations. Upstream and downstream variants encompass 5 kb genomic regions from the START and STOP sites, respectively. The Ensembl Variant Effect Predictor tool was used for SNP effect prediction using the Express617 reference genome (Lee *et. al.*, 2020). *Reflects the mean SNP counts from 497 IRFFA 4x pools.

**Supplementary Figure 3** - Characterization of “Non-EMS” type (A) Transitions (T→C and A →G) and (B) Transversions (C→A, G→T, A→C, T→G, T→A, A→T, G→C and C→G) with predicted effects on a genome-wide scale. Boxplots show the distribution of SNP effects as a mean of 497 4x pools. Mutation effects are predicted using filtered SNPs with DP≥10, AD=12.5%-60% and MQ≥30. The Ensembl Variant Effect Predictor release 99 was used in offline mode. Mutation effects were predicted within the gene models of the Express617 genome. All gene IDs were extracted from the general feature format (GFF) of the Express617 reference genome. Splice site variants include acceptor and donor site mutations. Upstream and downstream variants are located within a distance of 5 kb from the transcription START and STOP sites, respectively. DP: Read depth, AD: Allele depth and MQ: Mapping quality.

**Supplementary Figure 4** - Characterization of “Non-EMS” type (A) Transitions (T→C and A →G) and (B) Transversions (C→A, G→T, A→C, T→G, T→A, A→T, G→C and C→G) with predicted effects exclusively within coding regions. Boxplots show the distribution of SNP effects as a mean of 497 4x pools. Mutation effects are predicted using filtered SNPs with DP≥10, AD=12.5%-60% and MQ≥30. The Ensembl Variant Effect Predictor release 99 was used in offline mode. Mutation effects were predicted within the gene models of the Express617 genome. All gene IDs were extracted from the general feature format (GFF) of the Express617 reference genome. Splice site variants include acceptor and donor site mutations. Upstream and downstream variants are located within a distance of 5 kb from the transcription START and STOP sites, respectively. DP: Read depth, AD: Allele depth and MQ: Mapping quality.

**Supplementary Figure 5** - Distribution and density of filtered mutations per 1 Mb windows across all sequenced pools for chromosomes A01-A10 and C01-C09. The red and blue lines represent the running average of frequency distribution of filtered mutations for EMS type (C→T and G→A transitions) and non-EMS type mutations (all other nucleotide substitutions), respectively across all pools. Black dots reflect the corresponding mutation densities per 1 Mb bins for each of the sequenced pools. The x-axis represents the length of chromosomes in Mb and each interval corresponds to a 1 Mb window. The y axis shows the number of SNPs located within each window. To smoothen the curves, the General Additive Model (GAM) in the ggplot2 package of R was used.

**Supplementary Figure 6** - Screenshot of the Integrative Genomics Viewer (IGV) showing read alignments from the 7 selected fourfold pools harboring previously detected EMS mutations. Pool IDs are named with the prefix “IR” or “D”. (1) Tracks showing 41 bp regions with 20 bases up and downstream of the detected mutations. (2) Coverage tracks showing read alignments with horizontal bars representing individual reads. SNPs are marked in different colors. (3) Inset windows representing the count details for each of the called nucleotides (A: Adenine, C: Cytosine, G: Guanine, T: Thymine or N: Unknown) and their share from all mapped reads at that position.

**Supplementary Figure 7** - Screenshot of the Integrative Genomics Viewer (IGV) showing read alignments from 12 fourfold pools harboring EMS mutations detected within candidate genes – *BnGTR2, BnMYB28* and *BnCYP79F1*. Pool IDs are named with the prefix “IR” or “D”. (1) Tracks showing 41 bp regions with 20 bases up and downstream of the detected mutations. (2) Coverage tracks showing read alignments with horizontal bars representing individual reads. SNPs are marked in different colors. (3) Inset windows representing the count details for each of the called nucleotides (A: Adenine, C: Cytosine, G: Guanine, T: Thymine or N: Unknown) and their individual share from all mapped reads at that position.

**Supplementary Figure 8** - Sanger sequencing results from experiments validating the presence of EMS-induced mutations detected in 12 4x pools (Supplementary Table 3). Using genomic DNA from the four representative individuals of the selected pools (Supplementary Table 3), PCR amplicons encompassing detected mutants were Sanger sequenced. Genomic sequences of the candidate genes were extracted from the Express617 reference genome. Location and type of detected mutations were annotated for each of the candidate genes (marked in blue arrows). Sanger sequencing reads were mapped to the corresponding candidate gene sequences to check for the presence of the expected mutations. Validated mutations from M_2_ individuals are marked with orange arrows. 4x pools are named with the “D” prefix. The 4 M_2_ individuals corresponding to each of the pools have been named with the pool IDs followed by the “_1”, “_2”, “_3” and “_4” suffixes. Sequence analysis was done using the CLC Main Workbench 7.

