## Supplementary Figures for "Direct access to millions of mutations by Whole Genome Sequencing of an oilseed rape mutant population"

**Supplementary Figure 1**

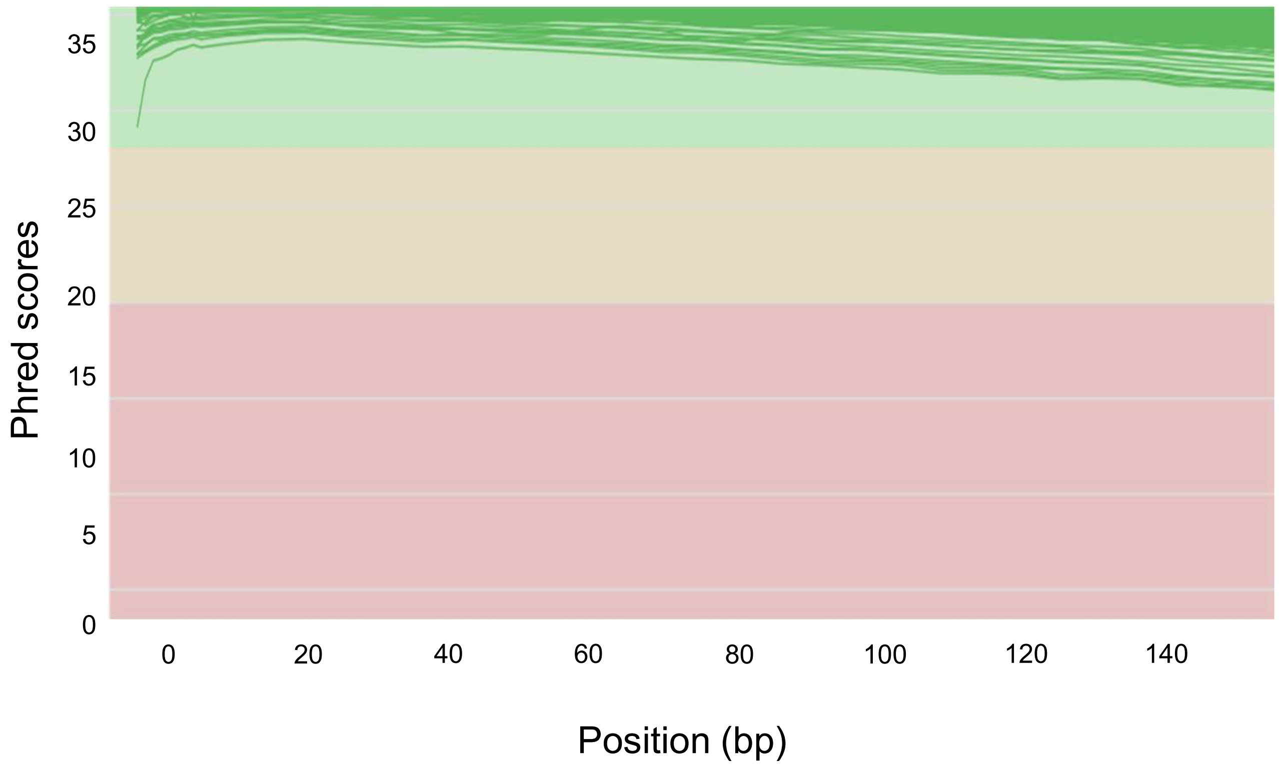

**Supplementary Figure 2**

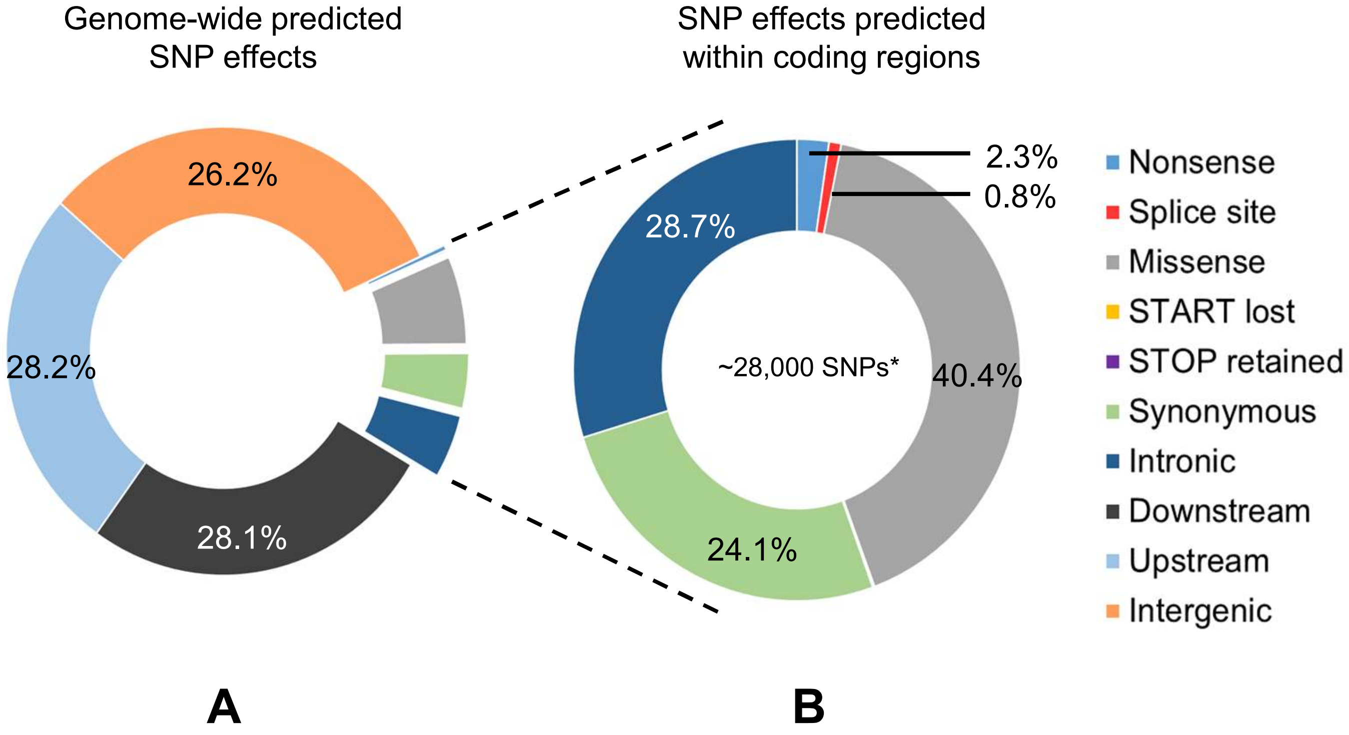

**Supplementary Figure 3**

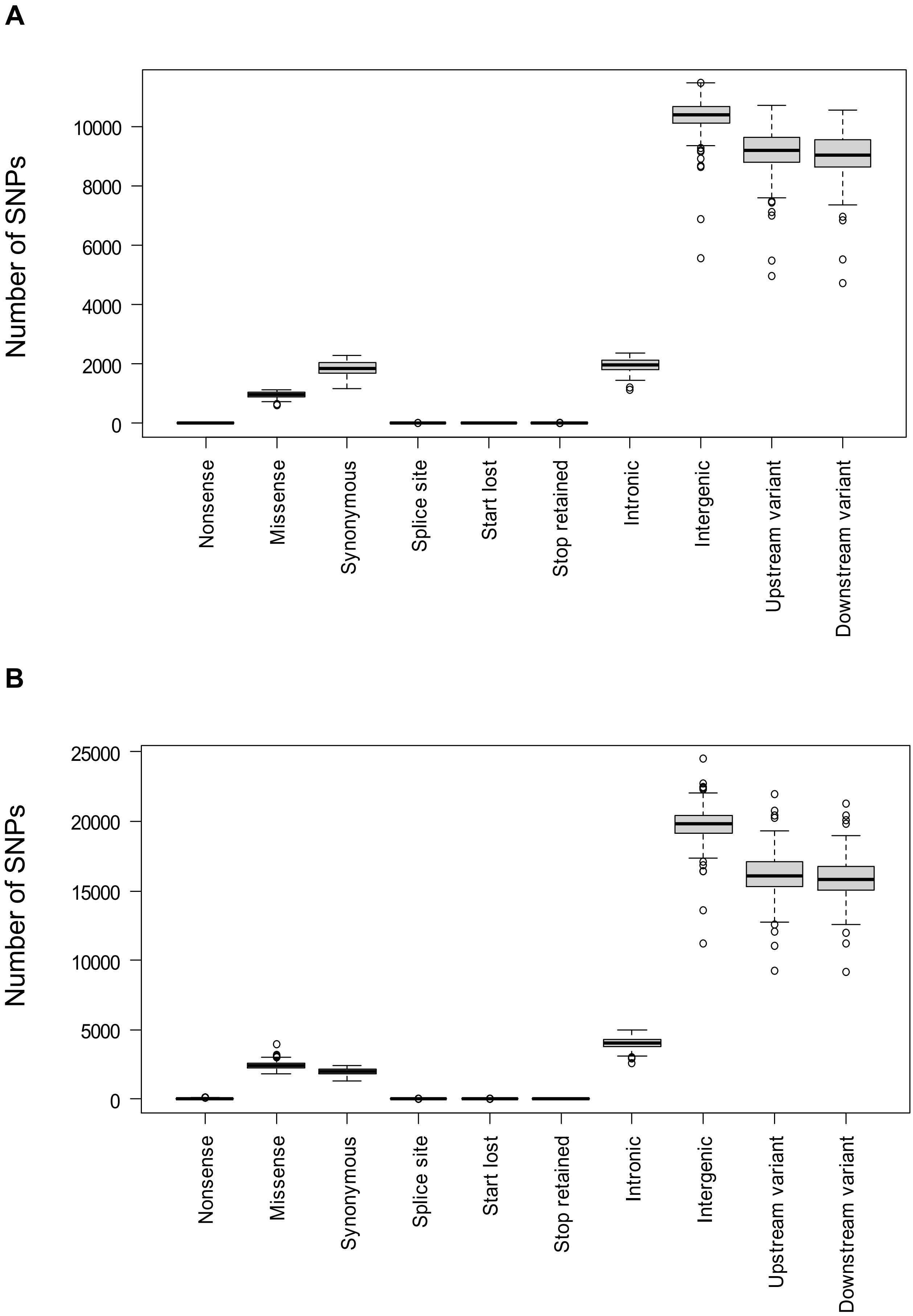

**Supplementary Figure 4**

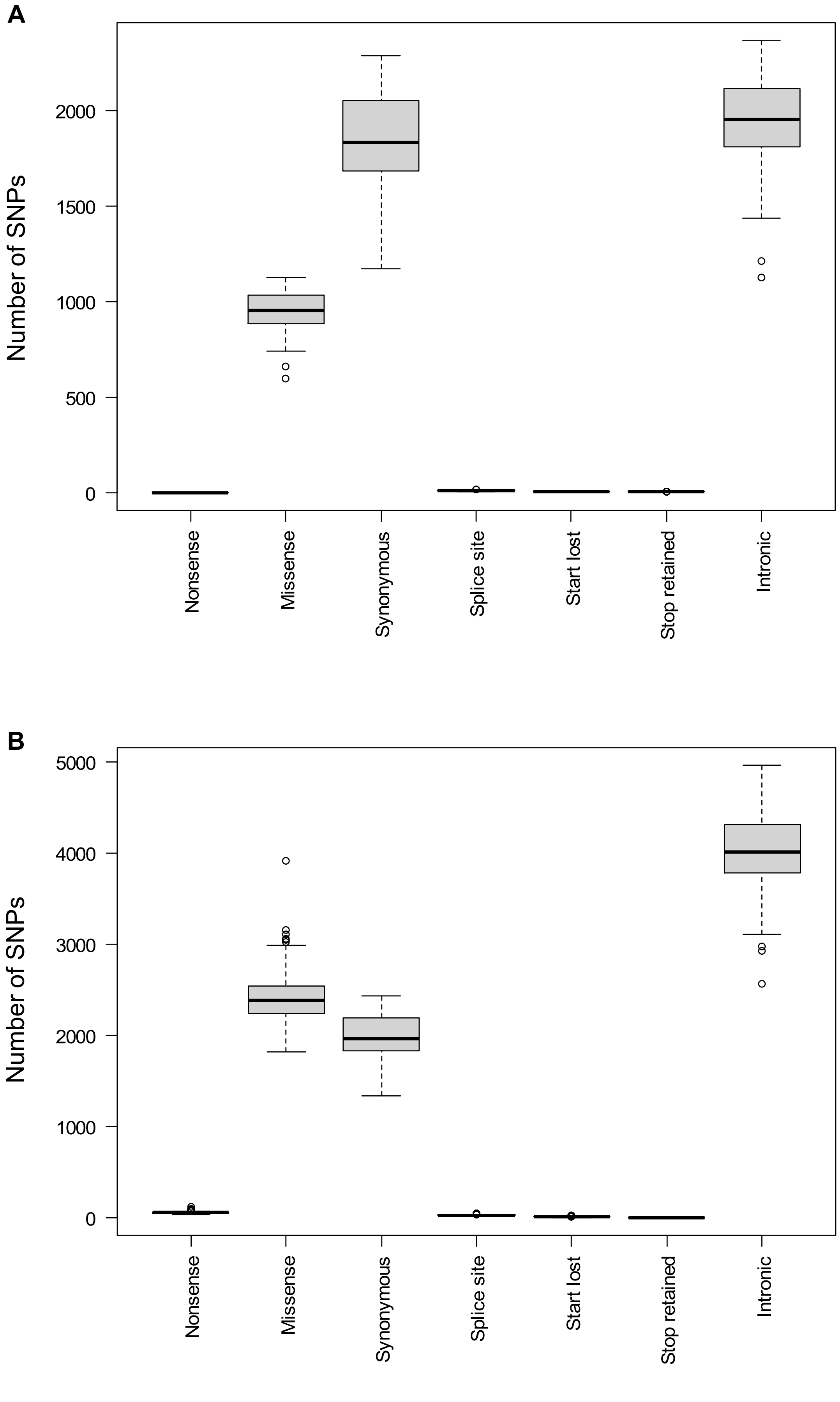

**Supplementary Figure 5**

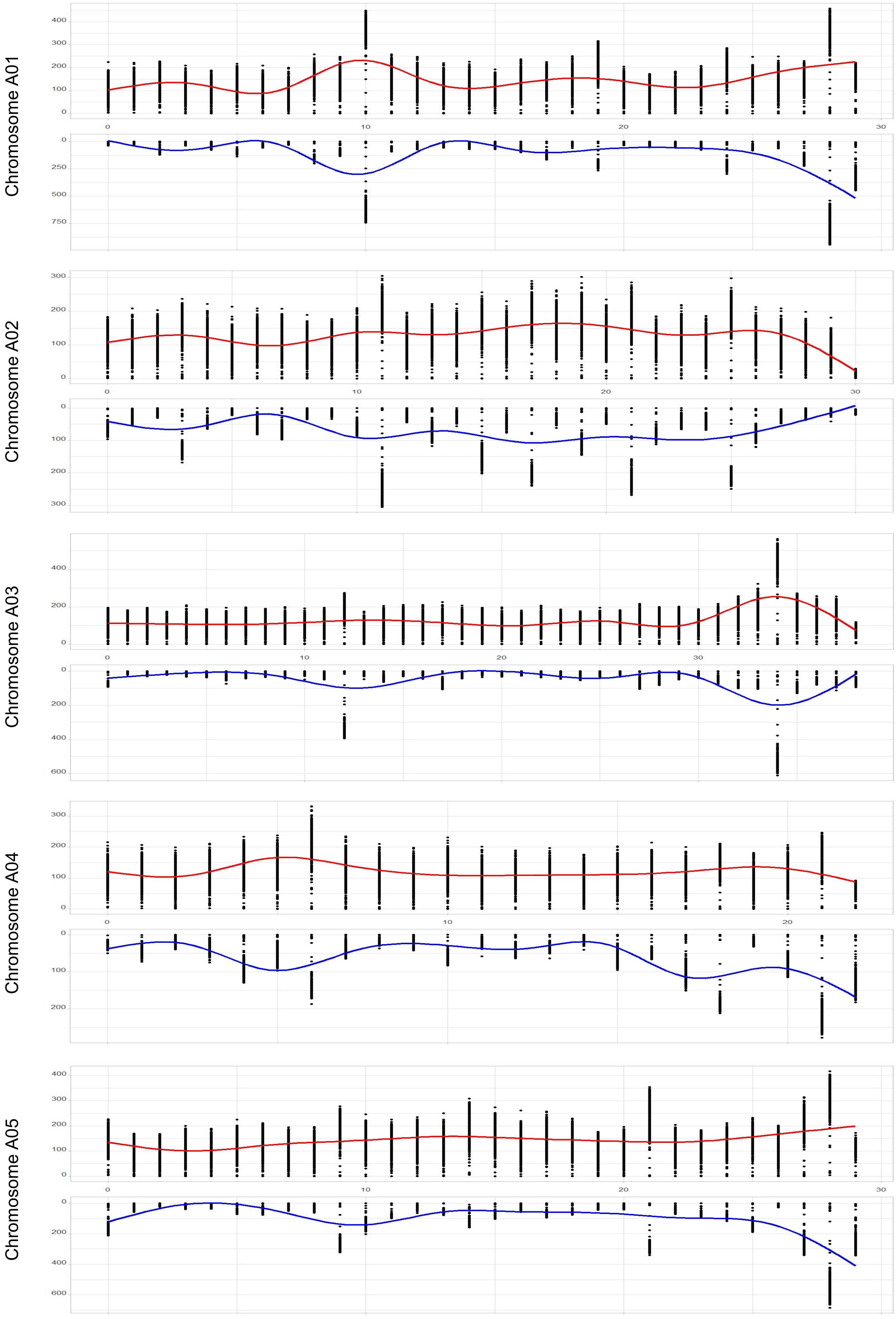

**Supplementary Figure 5 *contd.***

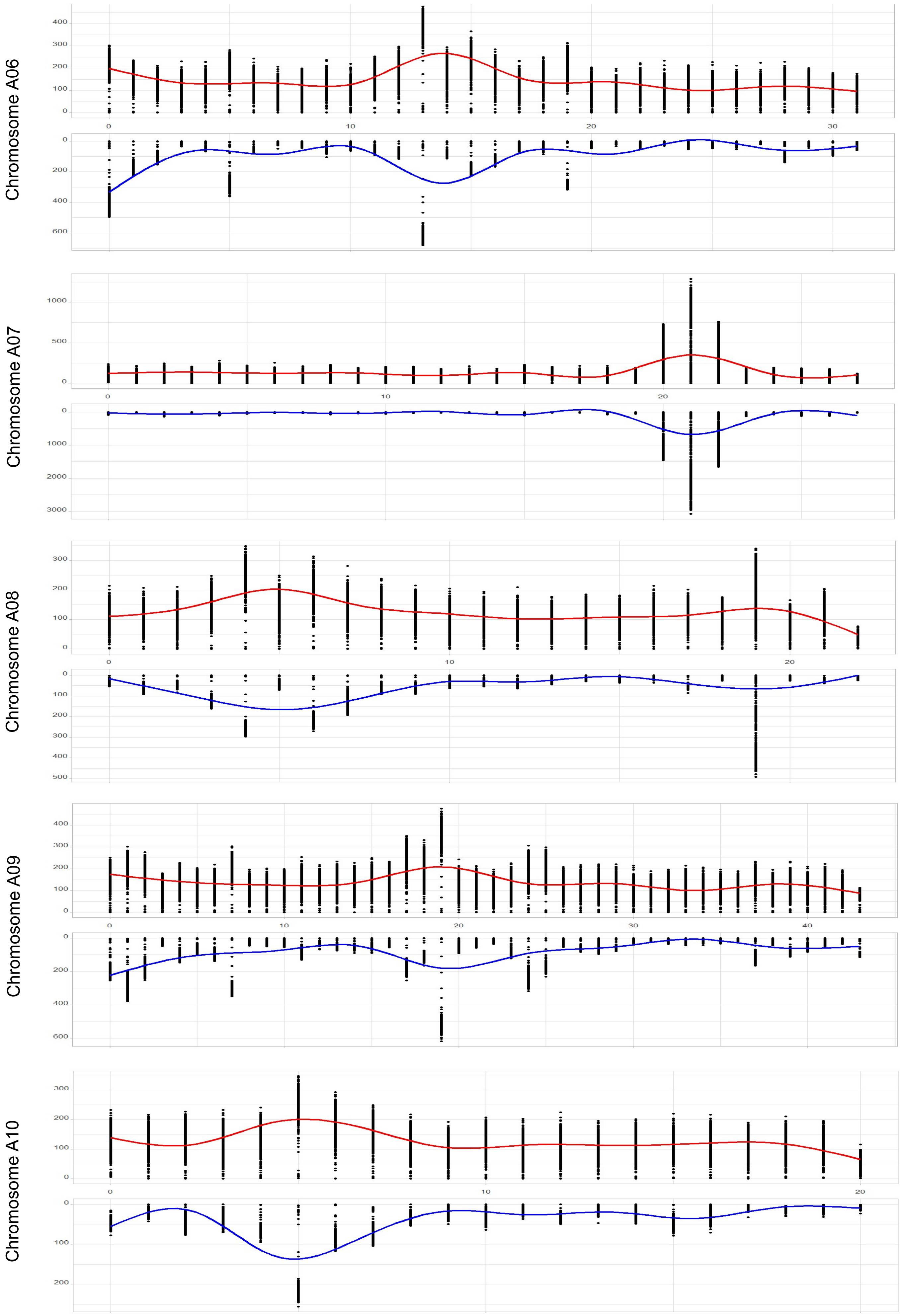

**Supplementary Figure 5 *contd.***

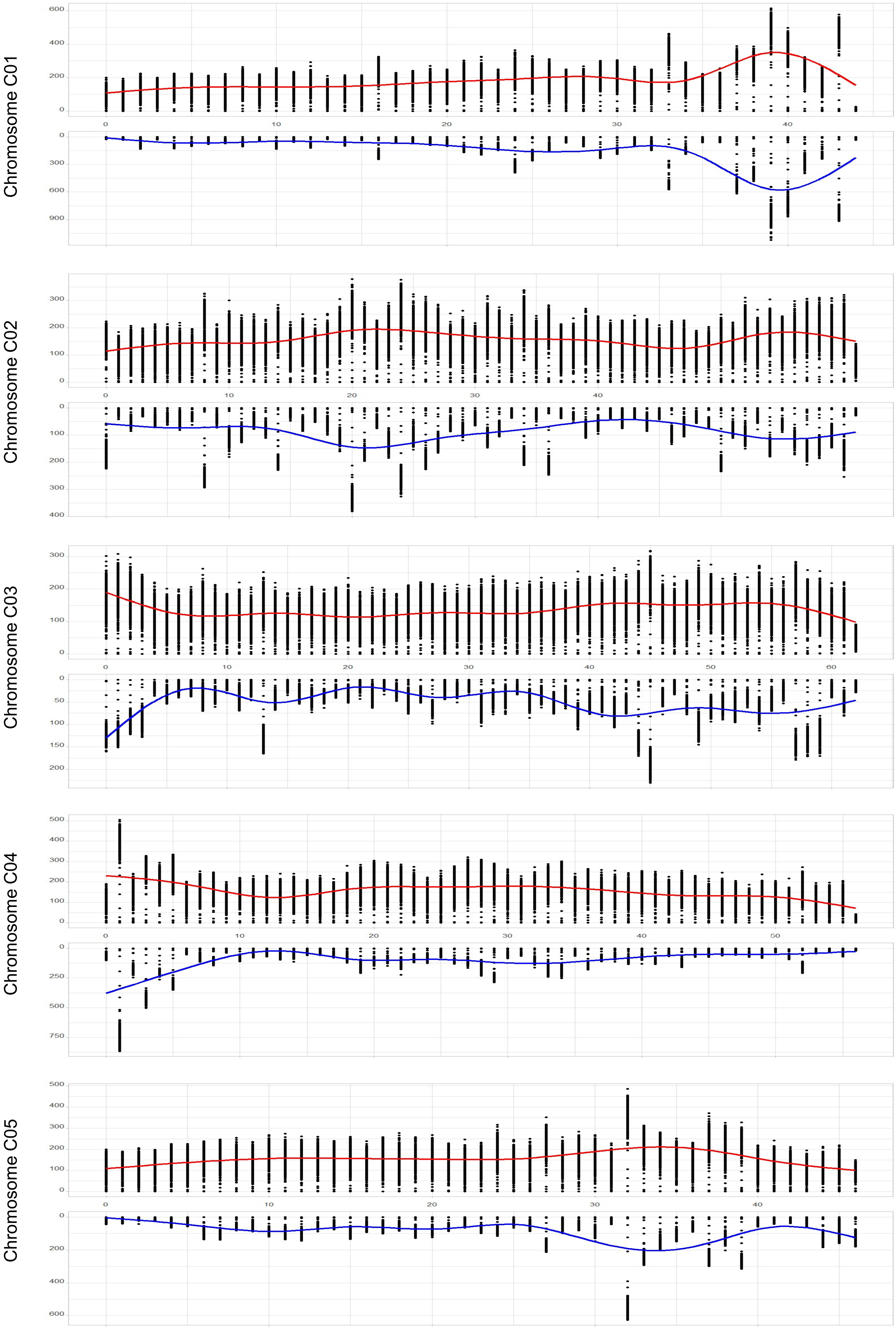

**Supplementary Figure 5 *contd.***

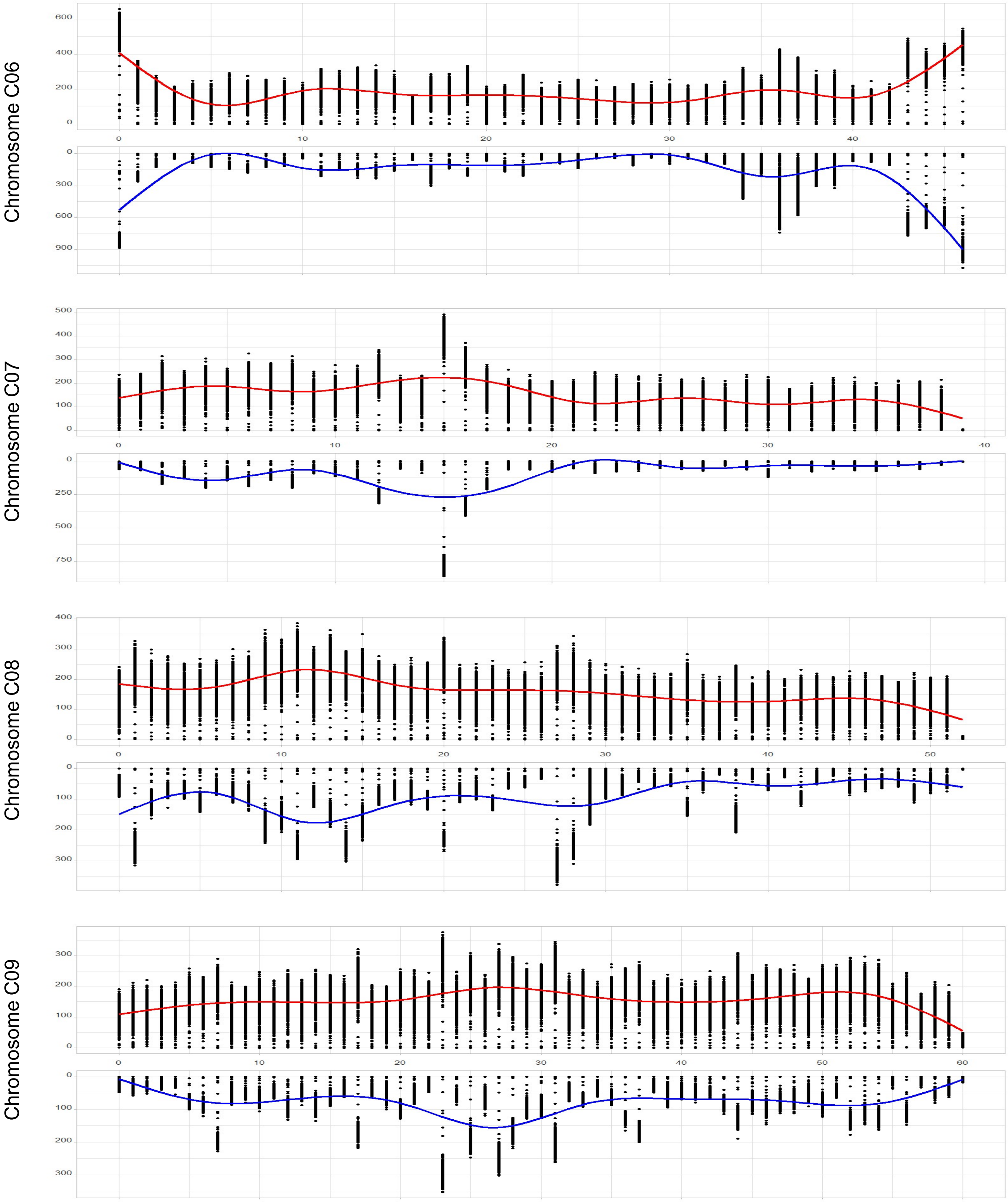

**Supplementary Figure 6**

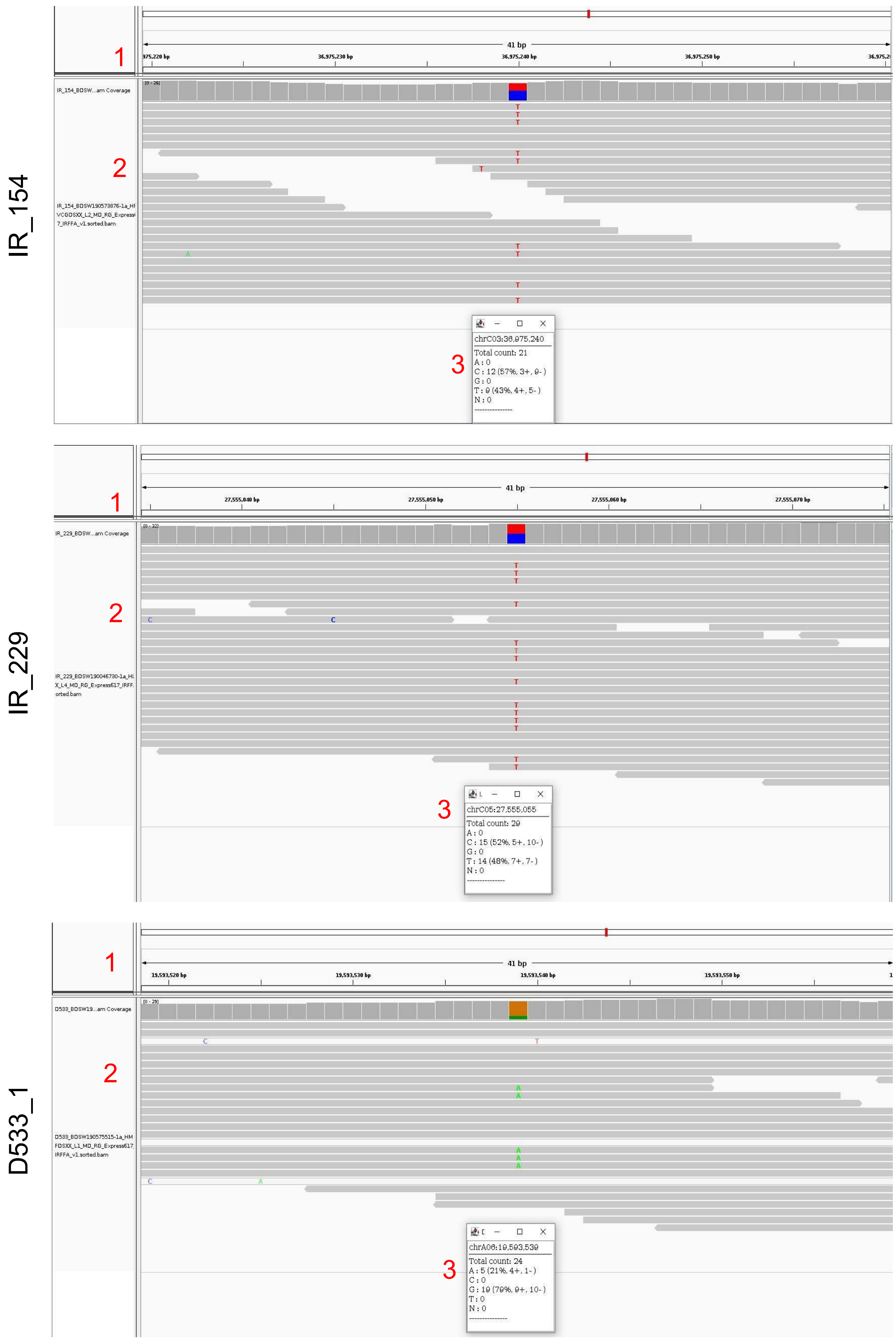

**Supplementary Figure 6 *contd.***

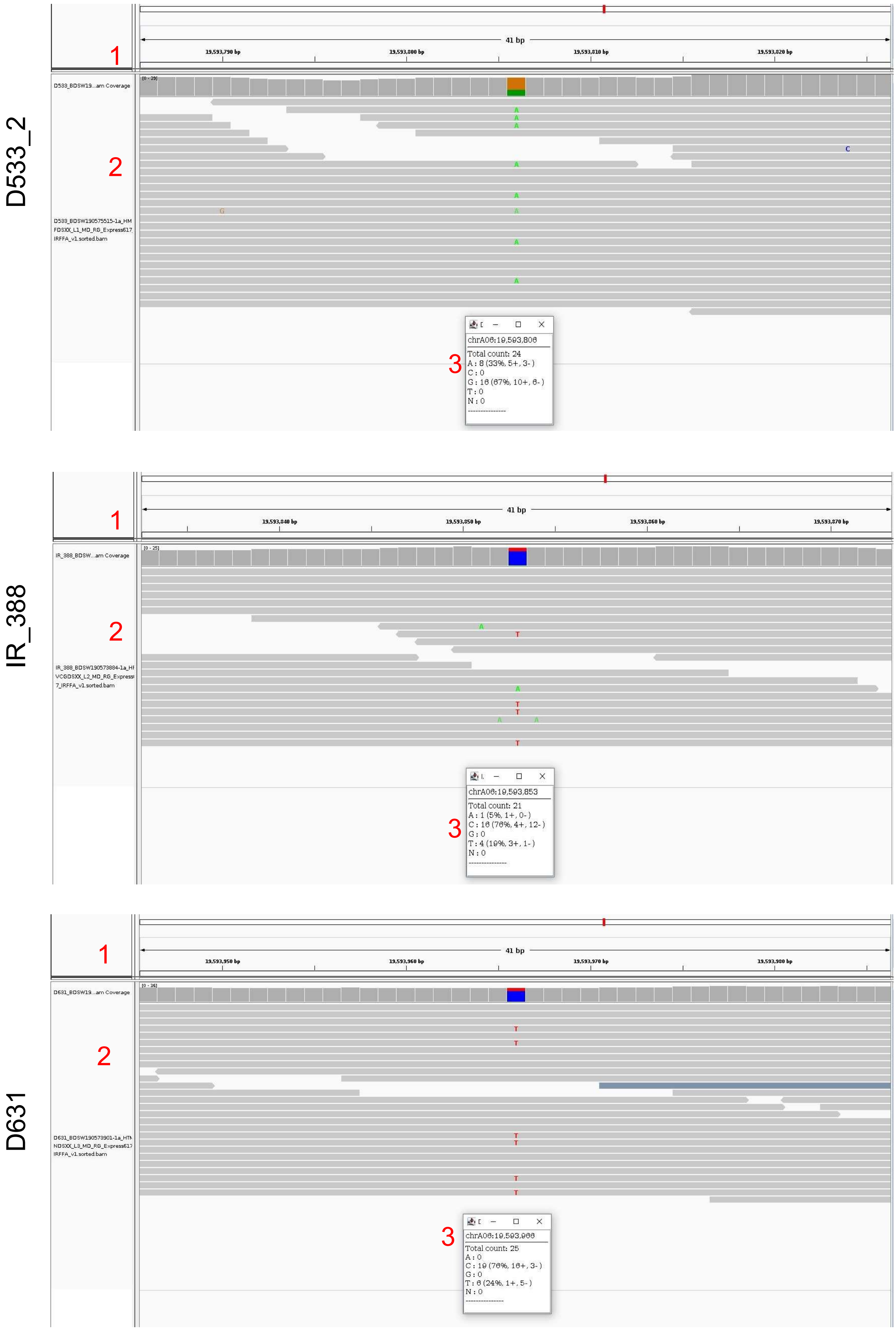

**Supplementary Figure 6 *contd.***

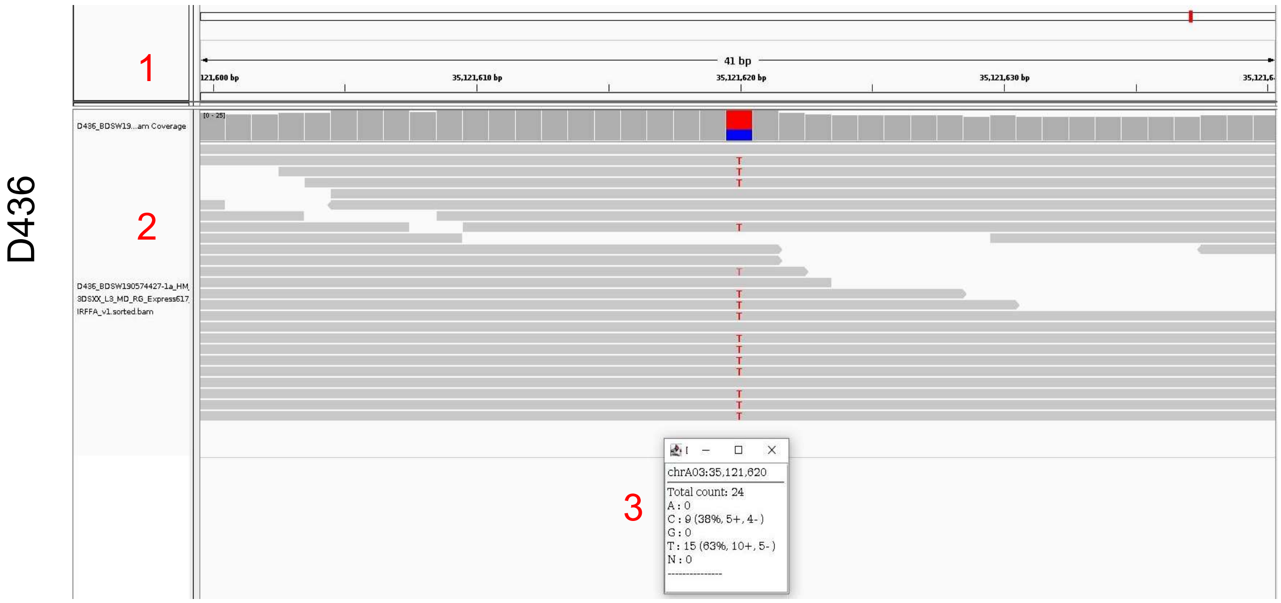

**Supplementary Figure 7**

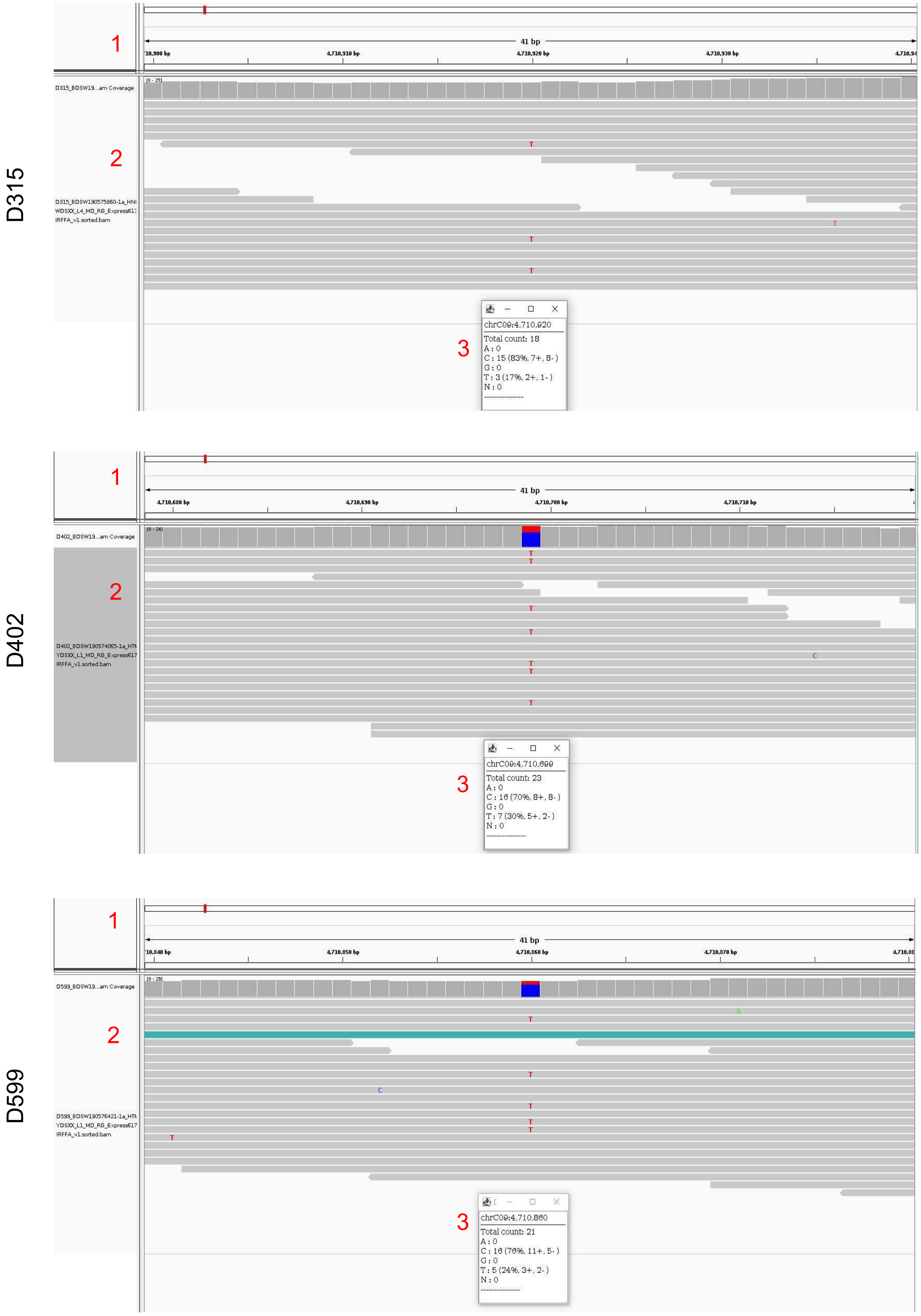

**Supplementary Figure 7 *contd.***

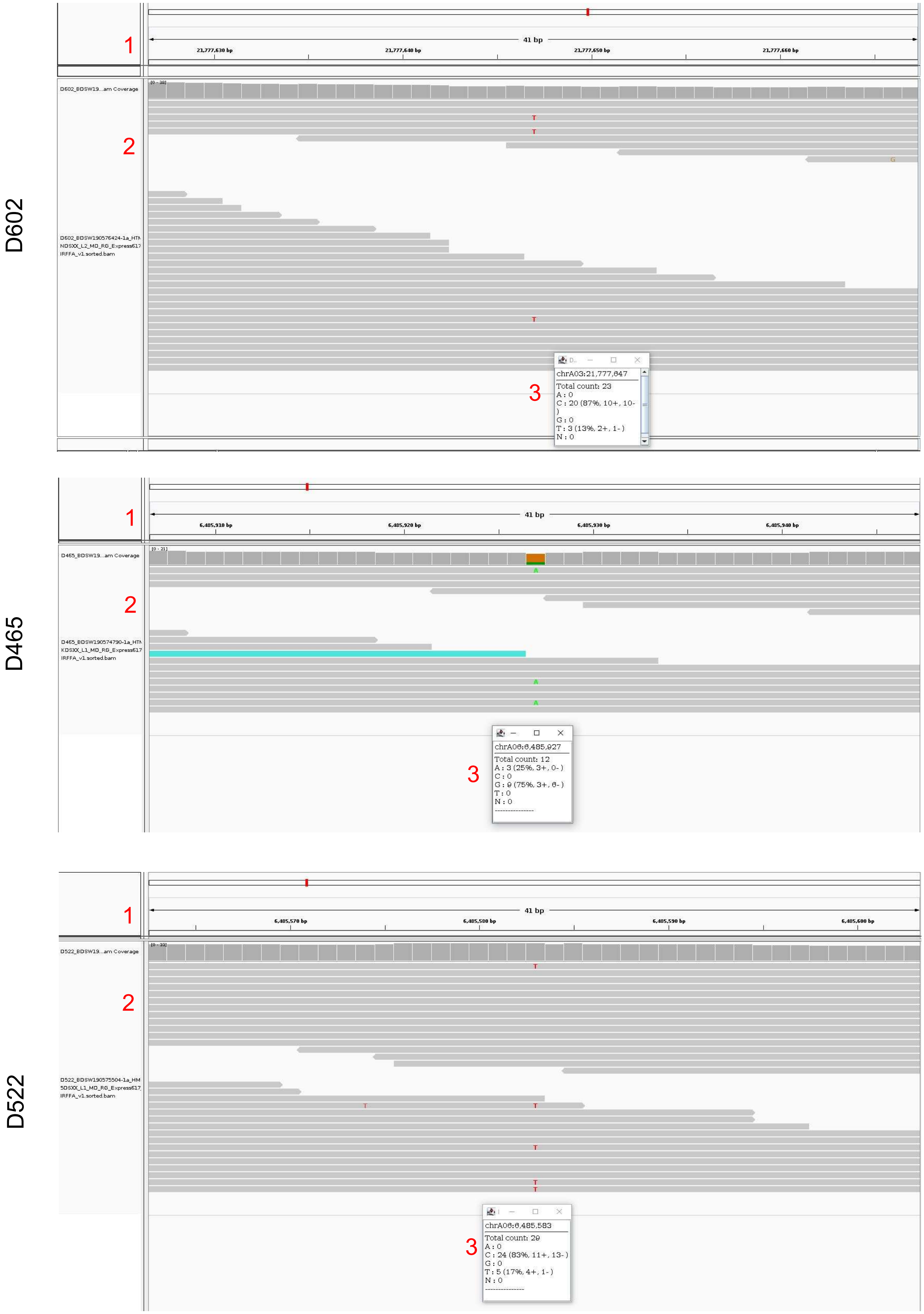

**Supplementary Figure 7 *contd.***

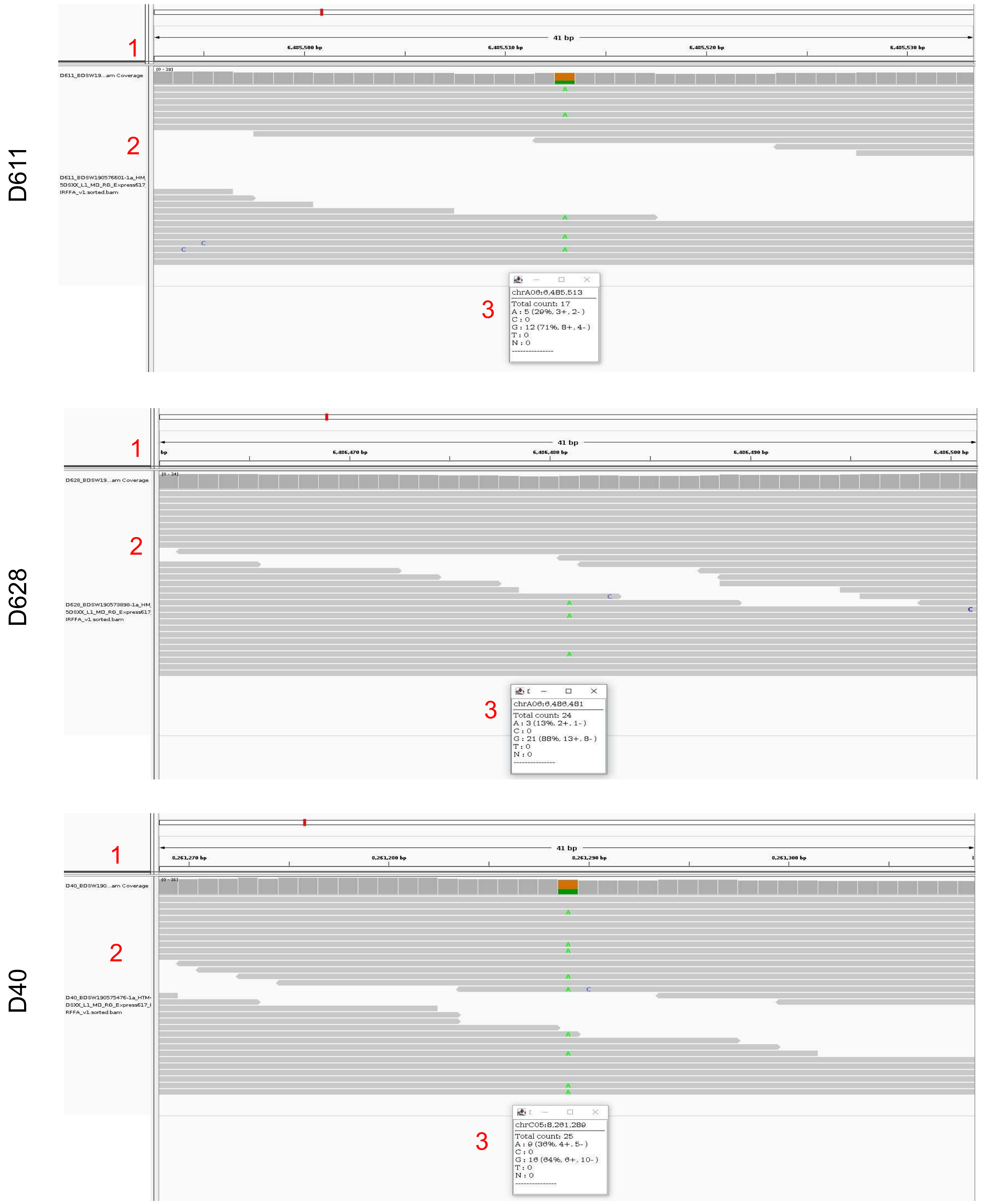

**Supplementary Figure 7 *contd.***

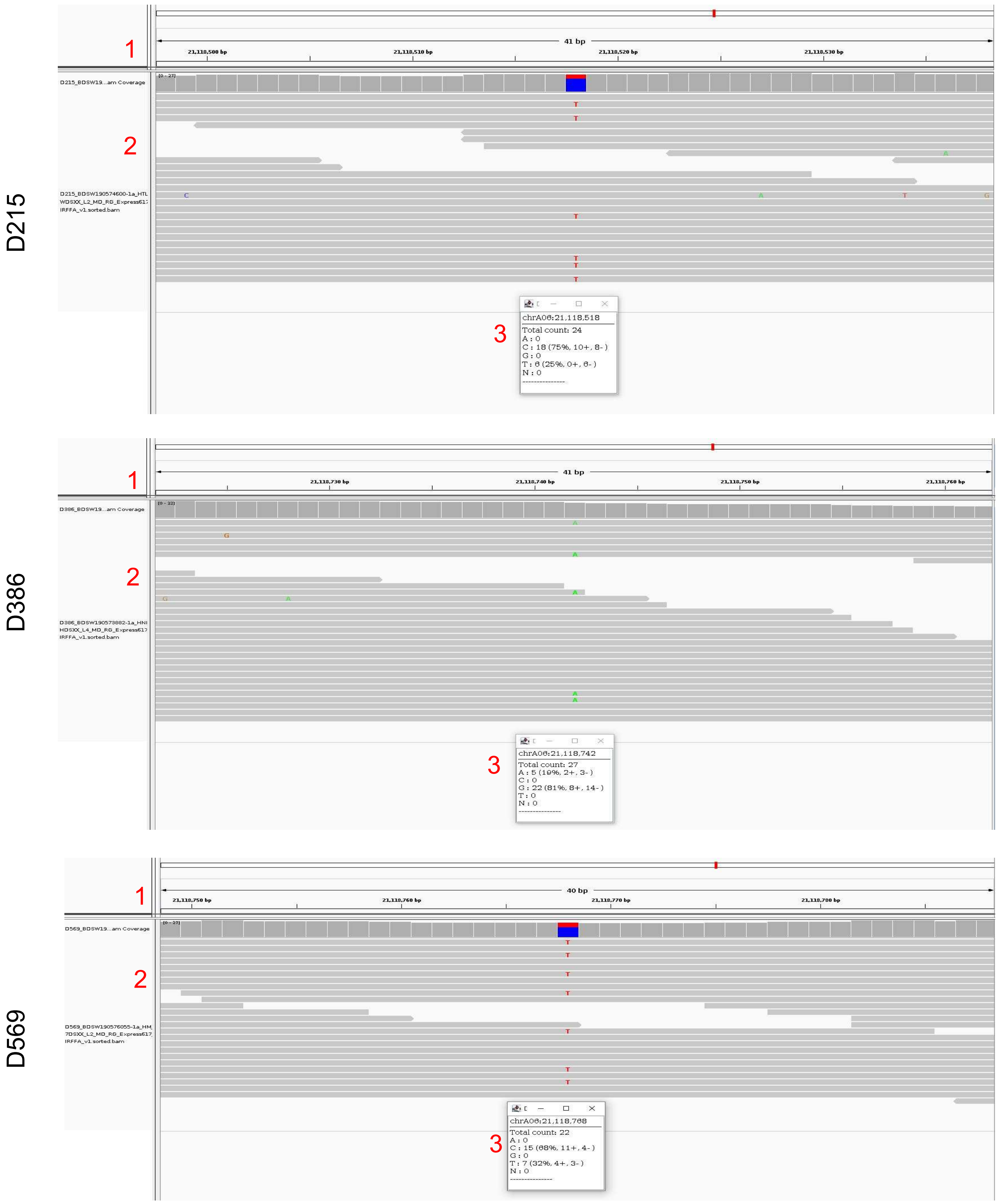

**Supplementary Figure 8**

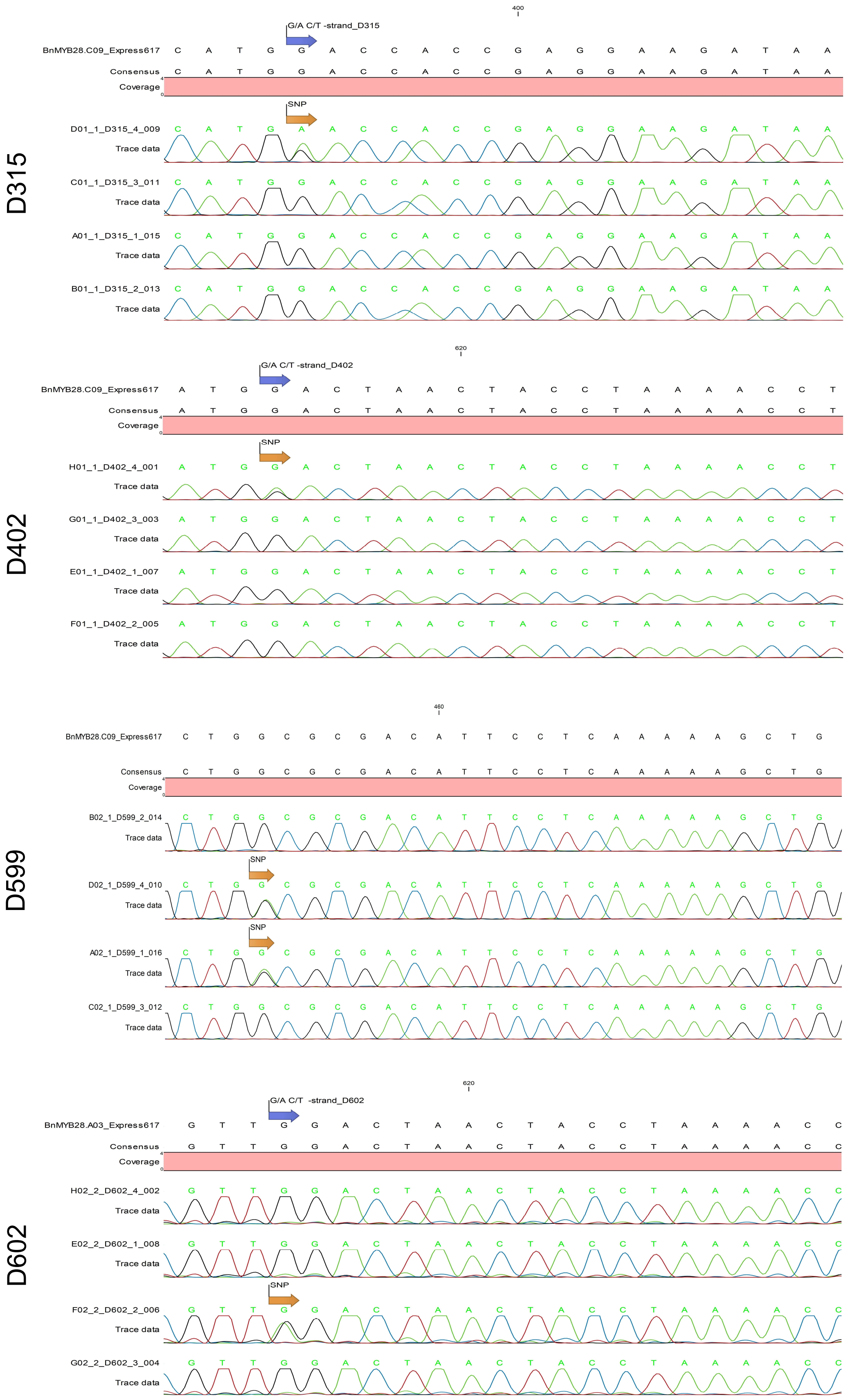

**Supplementary Figure 8 *contd.***

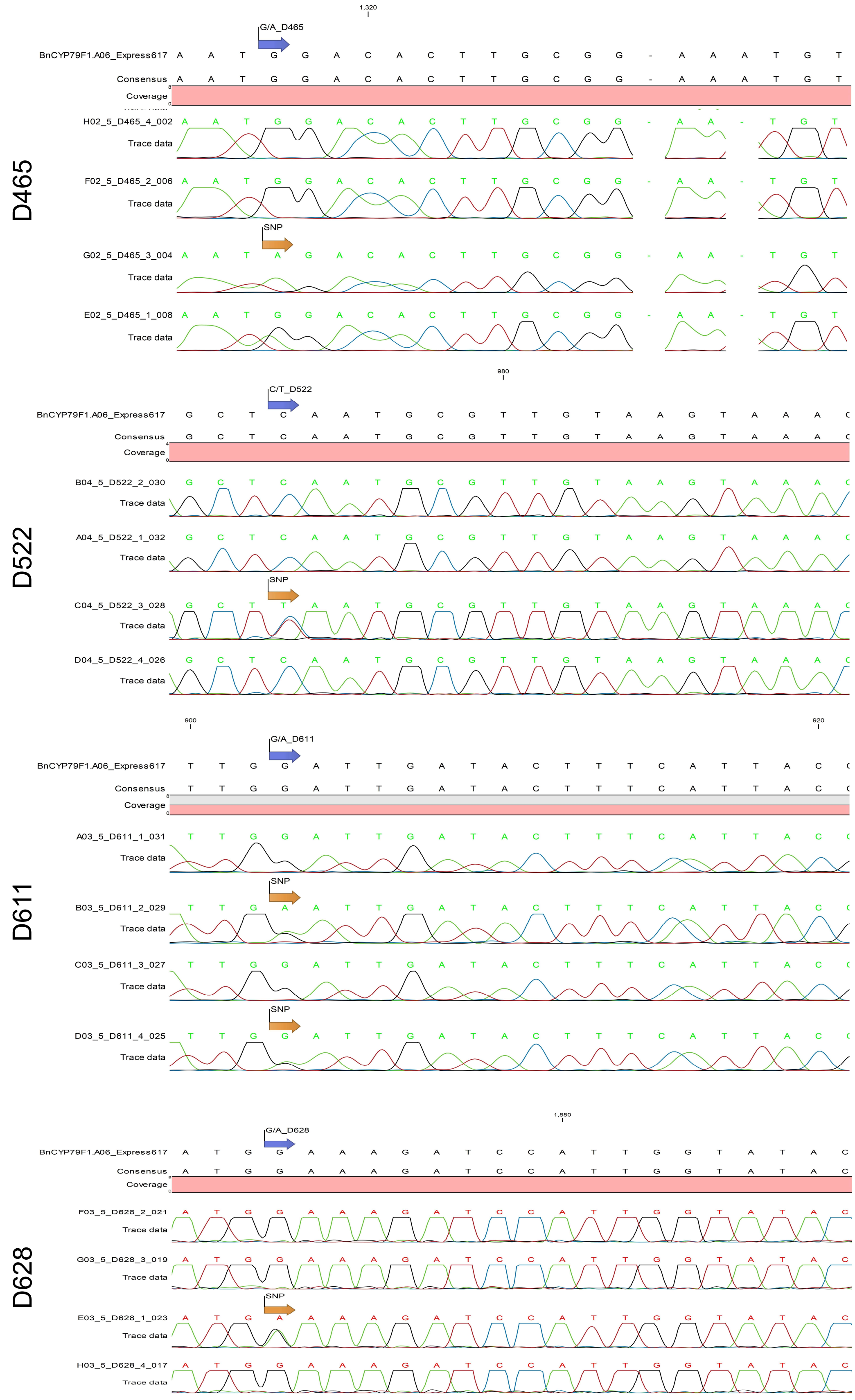

### Supplementary Figure 8 *contd.*

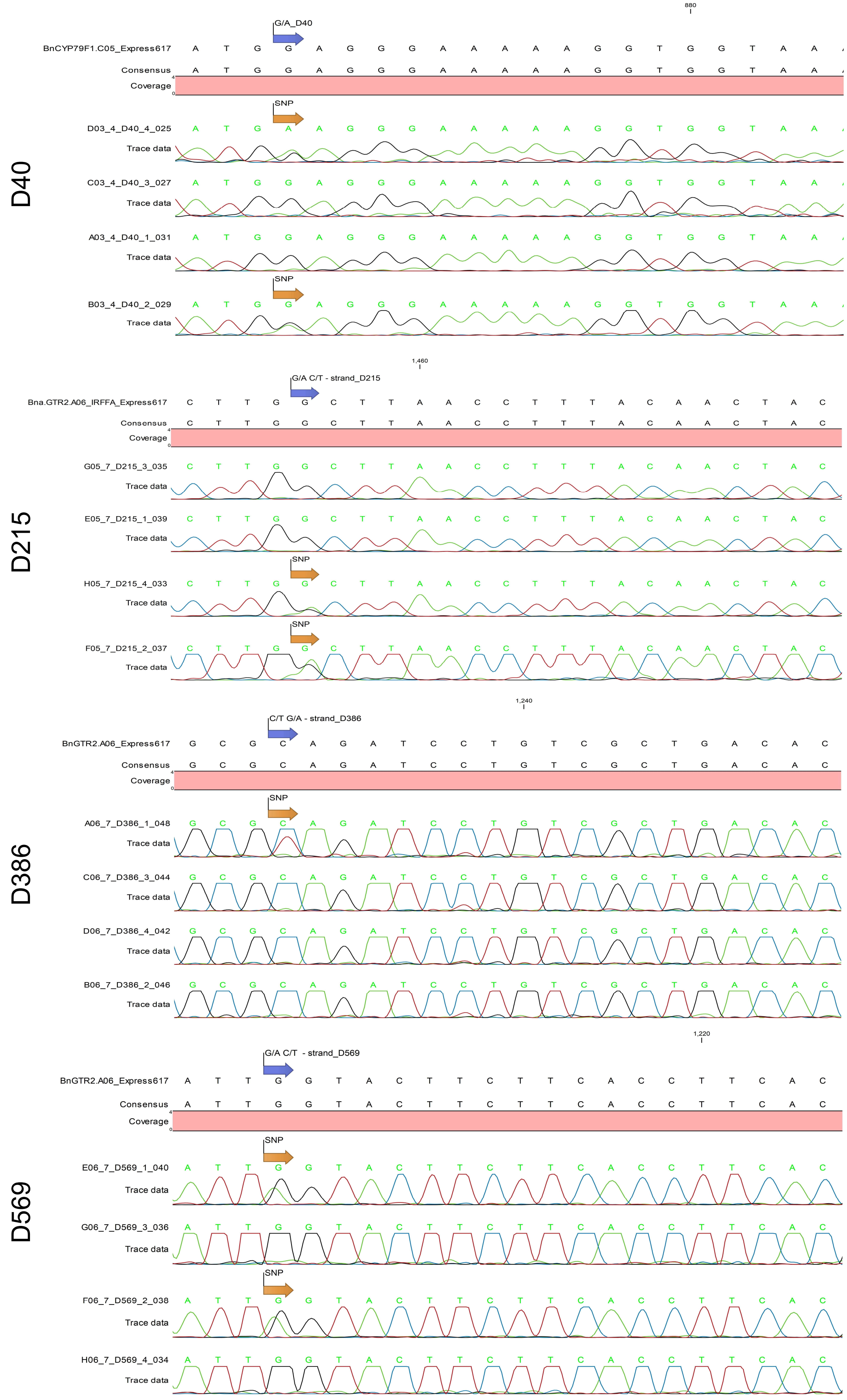
