## Supplementary Tables for "Direct access to millions of mutations by Whole Genome Sequencing of an oilseed rape mutant population"

### Supplementary data

#### Supplementary Tables

**Supplementary Table 1 –** Summary of EMS-type transitions originating from the annotated and non-annotated regions of the Express617 genome. Regions of the Express617 reference assembly without chromosomal annotations for the A- or C sub-genomes are denoted as non-annotated regions. Share of C→T and G→A transitions located within annotated and non-annotated regions was calculated as an average from 497 sequenced 4x pools.

| | **Length (bp)** | **Average SNPs /4x pool** | **%** | **Mutation frequency per Mb** |
|---|---|---|---|---|
| chrA01- chrA10 | 295,690,213 | 41,413 | 26.5 | 35 |
| chrC01- chrC09 | 468,939,566 | 76,819 | 49.1 | 41 |
| Non-annotated regions | 160,465,280 | 38,248 | 24.4 | 59 |
| **Total** | 925,095,059 | 156,480 | 100 | 45 |

Chromosome lengths are based on the assembled Express617 reference genome (Lee et al. 2020).

**Supplementary Table 2** - Summary of validation experiments to confirm mutations in 7 $M_2$ DNA pools harboring previously detected and characterized EMS mutations within 4 candidate gene families, *BnREF*, *BnSGT, BnSFAR4* and *BnSFAR1*. For each of the selected mutants, corresponding $M_2$ families were identified. Individual read alignments from each of the 4x pools (named with prefix "D" or "IR") representing the selected mutant $M_2$ families were visualized for regions harboring expected mutations.

| Gene family | Gene name | Darmor-bzh gene ID [1] | Mutation position [2] | Mutation effect | Mutation type | Pool ID | Express617 gene ID [3] | Mutation position in Express617 reference [2] | References |
|---|---|---|---|---|---|---|---|---|---|
| *BnREF* | *BnREF1.C03* | *BnaC03g43810D* | G 3329 A | D 392 N | Missense | IR_154 | *C03p045600* | C03_36975240 | Emrani et al. (2015) |
| *BnSGT* | *BnSGT.C05* | *BnaC05g30670D* | G 1054 A | E 352 K | Missense | IR_229 | *C05p028480* | C05_27555055 | Harloff et al. (2012) |
| *BnSFAR4* | *BnSFAR4.A06a* | *BnaA06g18900D* | G 497 A | G 166 E | Missense | D533 | *A06p021720* | A06_19593539 | (Karunarathna et al., 2020) |
| | *BnSFAR4.A06a* | | G 764 A | Splice site mutation | | D533 | | A06_19593806 | |
| | *BnSFAR4.A06a* | | C 811 T | P 231 L | Missense | IR_388 | | A06_19593853 | |
| | *BnSFAR4.A06a* | | C 924 T | Q 269 * | Nonsense | D631 | | A06_19593966 | |
| *BnSFAR1* | *BnSFAR1.Ann* | *Not annotated* | G 411 A | G 103 R | Missense | D436 | *A03p056350* | A03_35121620 | |

[1] Based on the Darmor-bzh reference genome (Genoscope)
[2] Relative to the translation START site
[3] Based on the Express617 reference genome (Lee et al., 2020)
*Premature stop codon mutation

**Supplementary Table 3 -** Summary of validation experiments to confirm the presence of mutations in 12 pools harboring EMS mutations within candidate gene families *BnMYB28*, *BnCYP79F1* and *BnGTR2*. 4x pools (named with prefix "D") with nonsense mutations for the candidate genes were first identified. Genomic DNA was isolated separately from the leaf samples of the 4 individuals (#1-4) bulked in each pool. Standard PCR with locus-specific primers was used to amplify regions encompassing the detected mutations. PCR fragments were Sanger sequenced to validate mutation presence.

| | | | | | **Flanking primers (5' → 3') [a]** | | **$M_2$ genotypes [b]** | | | | |
|---|---|---|---|---|---|---|---|---|---|---|---|
| **Gene name** | **Express617 gene ID [1]** | **Pool ID** | **Mutation position[1]** | **% Read coverage of mutation in 4x pool (AD/DP) [2]** | **Forward primer** | **Reverse primer** | **#1** | **#2** | **#3** | **#4** | **Observed coverage of mutant alleles (%) [3]** |
| *BnMYB28* | *C09p006910* | D315 | C 4710920 T | 3/18 = 16.7 | gagcttctctattctcatcctag | gaccgaccacctaaagaccag | $A_eA_e$ | $A_eA_e$ | $A_eA_e$ | $A_1A_e$ | 12.5 |
| | | D402 | C 4710699 T | 7/23 = 30.4 | | | $A_eA_e$ | $A_eA_e$ | $A_eA_e$ | $A_2A_e$ | 12.5 |
| | | D599 | C 4710860 T | 5/21 = 23.8 | | | $A_3A_e$ | $A_eA_e$ | $A_eA_e$ | $A_3A_e$ | 25.0 |
| | *A03p040980* | D602 | C 21777647 T | 3/23 = 13.0 | gcattcttgggtgtttttgaggg | gcgttggaactatcctcttc | $A_eA_e$ | $A_4A_e$ | $A_eA_e$ | $A_eA_e$ | 12.5 |
| *BnCYP79F1* | *A06p011120* | D465 | G 6485927 A | 3/12 = 25.0 | cattgacgagagggtggagc | ccatcatgatcgtcccgactttg | $B_eB_e$ | $B_eB_e$ | $B_1B_e$ | $B_eB_e$ | 12.5 |
| | | D522 | C 6485583 T | 5/29 = 17.2 | | | $B_eB_e$ | $B_eB_e$ | $B_2B_e$ | $B_eB_e$ | 12.5 |
| | | D611 | G 6485513 A | 5/17 = 29.4 | | | $B_eB_e$ | $B_3B_e$ | $B_eB_e$ | $B_3B_e$ | 25.0 |
| | | D628 | G 6486481 A | 3/24 = 12.5 | | | $B_4B_e$ | $B_eB_e$ | $B_eB_e$ | $B_eB_e$ | 12.5 |
| | *C05p012900* | D40 | G 8261289 A | 9/16 = 56.3 | ccgacttcgcagaccggcct | cggcctattccaggacgg | $B_eB_e$ | $B_4B_e$ | $B_eB_e$ | $B_4B_e$ | 25.0 |
| *BnGTR2* | *A06p023060* | D215 | C 21118518 T | 6/24 = 25.0 | cggtcgatacaagactctaactg | cttcttaaggtaggcacgagcac | $C_eC_e$ | $C_1C_e$ | $C_eC_e$ | $C_1C_e$ | 25.0 |
| | | D386 | G 21118742 A | 5/27 = 18.5 | | | $C_2C_e$ | $C_eC_e$ | $C_eC_e$ | $C_eC_e$ | 12.5 |
| | | D569 | C 21118768 T | 7/22 = 31.8 | | | $C_3C_e$ | $C_3C_e$ | $C_eC_e$ | $C_eC_e$ | 25.0 |

[a] Primers used to amplify the detected mutants from the Express617 TILLING population via standard PCR.

[b] Alleles observed after Sanger sequencing of PCR products from the four segregating $M_2$ individuals constituting the 4x pools.
$A_e$, $B_e$ and $C_e$ are Express617 wild type alleles for the respective paralogs of *BnMYB28*, *BnCYP79F1* and *BnGTR2*, respectively. Mutant alleles (are represented by suffix 1-4 ( $A_1$-$A_4$, $B_1$-$B_4$ and $C_1$-$C_3$.
[1] Express617 reference genome (Lee et al., 2020).
[2] Share of reads possessing mutations (AD) out of all reads mapped at that position (DP).
[3] Share of mutant alleles in the 4 $M_2$ individuals observed after Sanger sequencing of PCR fragments.
